# Positional Scanning and Computational Modeling Reveal Determinants of Legumain Transpeptidase Activity

**DOI:** 10.64898/2025.12.12.693912

**Authors:** Rupert Klaushofer, Sven O. Dahms, Hans Brandstetter, Elfriede Dall

## Abstract

Legumains are cysteine proteases that, in addition to their canonical hydrolase function, can act as peptide ligases or transpeptidases. In humans, this activity becomes particularly relevant under pathophysiological conditions, where legumain relocalizes to near-neutral pH compartments favoring ligation/transpeptidation over hydrolysis. Here, we combined *in vitro* positional scanning with *in silico* substrate profiling to elucidate the substrate determinants governing human legumain-mediated peptide cyclization. We identified glycine residues at P1″ and P1′ and basic residues at P2′/P2″ as key determinants of human legumain-mediated peptide cyclization. Guided by these insights, we designed an optimized substrate exhibiting substantially enhanced cyclization efficiency. Computational analysis not only recapitulated the experimental observations but also predicted a covalent inhibition mechanism involving a P1′ cysteine, revealed a *k*_cat_-tuning switch embedded within the substrate, and highlighted its potential for developing high-performance fluorogenic substrates. Collectively, these findings advance the mechanistic understanding of legumain’s transpeptidase activity and provide a framework for developing selective probes and inhibitors across the legumain family and related cysteine proteases.

**GRAPHICAL ABSTRACT:** 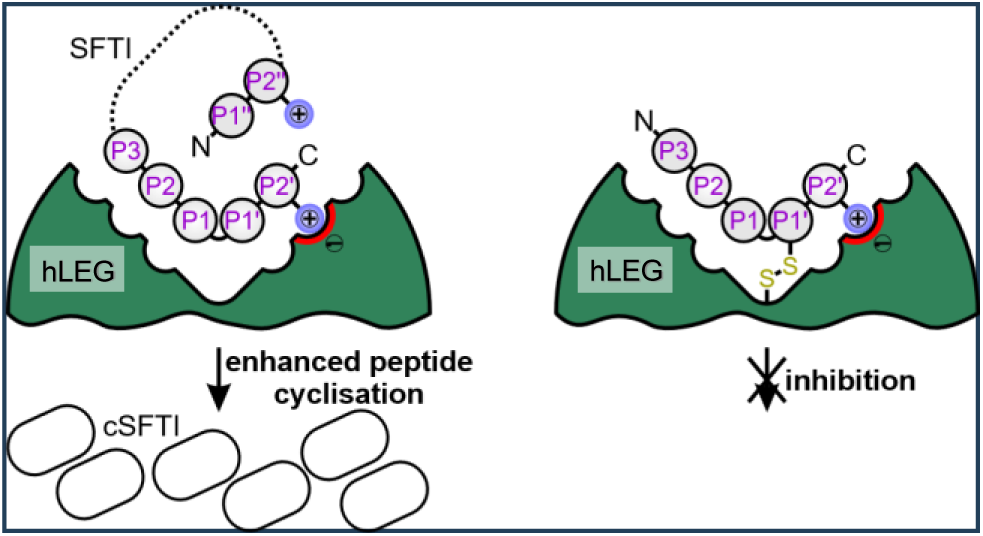

## INTRODUCTION

Legumains are cysteine proteases found throughout eukaryotes, where they perform diverse biological functions ranging from protein turnover in endo-lysosomal compartments to post-translational peptide cyclization in plants and certain invertebrates ^1–4^. In humans, legumain is primarily localized to the endo-lysosomal system, where it contributes to the processing of antigens for presentation on MHC class II complexes ^5–7^. In a pathophysiological context, legumain is overexpressed in the majority of human solid tumors, including breast cancer and colorectal cancer ^8–10^. Elevated expression is associated with enhanced tissue invasion and metastasis, and consequently correlates with poor clinical prognosis ^11–13^. Under these pathological conditions, legumain has been detected not only in lysosomes but also translocated to the nucleus, cytoplasm, and extracellular space [16, 18-21]. Recent evidence further suggests that similar mislocalization occurs in the aged brain, where legumain facilitates the aggregation of proteins implicated in neurodegenerative diseases, ultimately contributing to neuronal damage ^14–17^. Collectively, these findings have positioned legumain as a promising therapeutic target in both cancer and neurodegeneration.

A defining feature of legumain is its strict requirement for Asn (or to a lesser extent, Asp) at the P1 position leading to its synonymous naming as the asparaginyl endopeptidase (AEP) ^18–21^. Beyond its canonical proteolytic function, certain legumain isoforms also act as peptide ligases or transpeptidases, making them of particular biotechnological interest, especially for the enzymatic synthesis of cyclic peptides in drug development ^3, 22–26^. Protease and ligase activities exist in a pH-dependent equilibrium: hydrolysis dominates under acidic conditions, whereas ligation and transpeptidation are favored at near neutral pH. Transpeptidation involves cleavage of an existing peptide bond coupled to the formation of a new one, typically when the nucleophilic amino group of an incoming peptide attacks an acyl-enzyme intermediate, substituting for water in the hydrolysis reaction. In contrast, ligation refers to the direct enzymatic or chemical joining of two free peptide termini to create a new peptide bond, without prior bond cleavage.

In plants, legumain transpeptidase activity drives the biosynthesis of a wide range of cyclic peptides, whereas in animals it has been less explored. Notably, the transpeptidase activity of human legumain (hLEG) may become particularly relevant under pathological conditions that lead to its relocalization to near-neutral compartments, environments conducive to peptide transpeptidation. In these contexts, precise understanding of substrate specificity is critical for designing efficient and selective transpeptidase substrates and inhibitors. However, the substrate preferences governing the transpeptidase activity of hLEG remain poorly characterized.

Here, we combine an *in vitro* positional peptide scanning screen using the sunflower trypsin inhibitor (SFTI) scaffold with *in silico* substrate profiling using AlphaFold to elucidate the substrate specificity of hLEG and the plant isoform *Arabidopsis thaliana* legumain β (AtLEGβ). This integrated approach enabled the development of optimized legumain cyclization substrates and provided mechanistic insights into catalysis and can guide the design of high-performance fluorogenic substrates.

## RESULTS & DISCUSSION

### 1. Design of peptide library

We previously established a peptide cyclization assay for hLEG using the sunflower trypsin inhibitor (SFTI) precursor peptide as substrate ^23^. In brief, the SFTI precursor is incubated with hLEG at pH 6.0, and product formation is monitored by MALDI-TOF MS (Fig. 1). Two products are observed: (i) a truncated linear peptide (L-SFTI), generated by hydrolysis of the prime-side tripeptide (P1’-P3’) Gly^15^-Leu^16^-Ala^17^, and (ii) a cyclic peptide (C-SFTI), which arises either by ligation of L-SFTI or directly from the precursor via transpeptidation.

**Figure 1.**
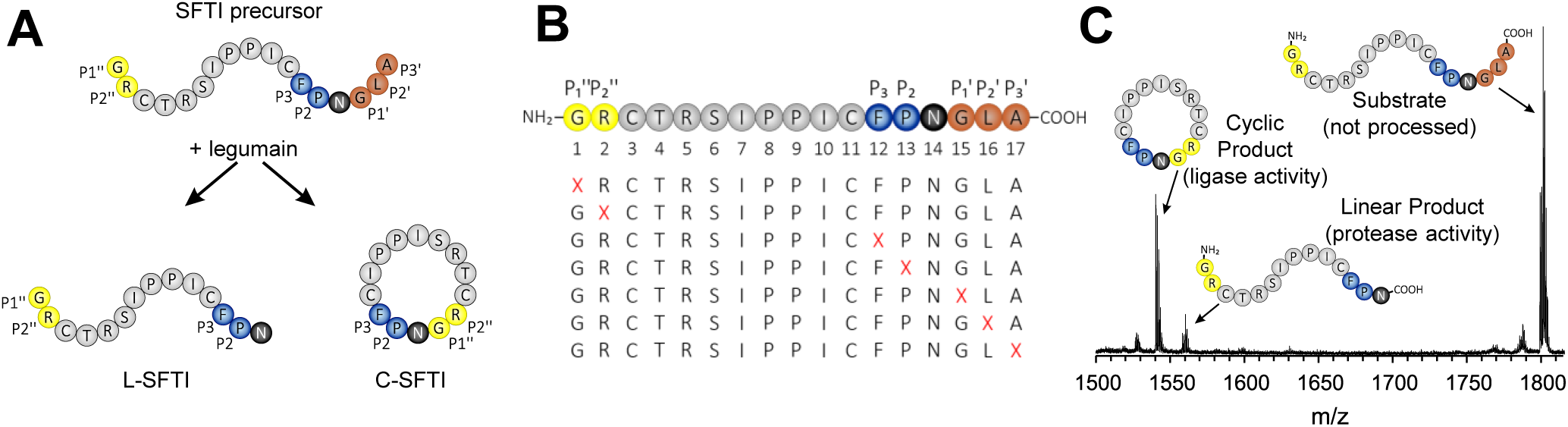
Design of the positional scanning assay. (A) The SFTI precursor peptide (top) is incubated with human legumain (hLEG), resulting in the formation of either a linear (L-SFTI) or a cyclic (C-SFTI) product. (B) A positional scanning peptide library was generated in which positions P2″, P1″, P3, P2, P1′, P2′, and P3′ were systematically substituted with the 20 proteinogenic amino acids. (C) Reaction products were analyzed by MALDI-ToF mass spectrometry.

To probe hLEG transpeptidase substrate specificity, we used the SFTI precursor as a scaffold for positional scanning. Based on our prior observation that a P1 Asn improves cyclization efficiency, we substituted the SFTI P1 Asp14 with Asn14 ^23^. We then focused on positions likely to contact the catalytic site. The SFTI-GL precursor contains a disulfide bond between Cys11 and Cys3; these residues represent the P4 and P3’’ positions, respectively, leading us to vary the positions limited by this disulfide bond. Specifically, positions P3, P2, P1’, P2’, P3’, P1’’, and P2’’ were individually substituted with each of the 20 proteinogenic amino acids (Fig. 1B). The P1 position was not varied, as asparagine is essential for hLEG to process the substrate. We synthesized a crude library of 140 peptides and incubated each sequence with hLEG at pH 6.0. Conversion to L-SFTI and C-SFTI was quantified by MALDI-TOF MS based on relative peak areas of the precursor, linear, and cyclic species (Fig. 1C).

### 2. Human legumain transpeptidase prefers basic residues at position P2’ and P2

Cyclic product yields were normalized to the alanine variant at each position and visualized in a heatmap (Fig. 2A). Notably, at P1’’, only glycine supported efficient cyclization. All other substitutions yielded little or no cyclic product, indicating that P1’’-Gly is essential for legumain-mediated transpeptidation (Fig. 2A,B). At P2’’, positively charged residues Arg, Lys and His enhanced cyclization, with alanine also being tolerated, whereas most branched amino acids were poorly accepted. At P3, phenylalanine strongly promoted the formation of cyclic product. As Phe is present in the native sequence, no improvement could be achieved upon variation at this position. At P2, the native proline was favorable, but substituting it with glycine further increased cyclization efficiency by a factor of 2.6. At P1’, glycine again proved optimal, likely due to its minimal steric hindrance and conformational adaptability, which can enhance cyclization efficiency. Variants containing Cys, His, Lys, or Ser at P1’ produced approximately half the amount of cyclic product relative to the Gly variant, while other substitutions were poorly tolerated. At P2’, basic residues (Lys, Arg) substantially increased cyclization efficiency; this preference mirrored that seen for P2’’, albeit much more pronounced. At P3’, residues such as Cys, His, Lys, and Arg enhanced product formation, although this position displayed lower selectivity overall compared to the others.

**Figure 2.**
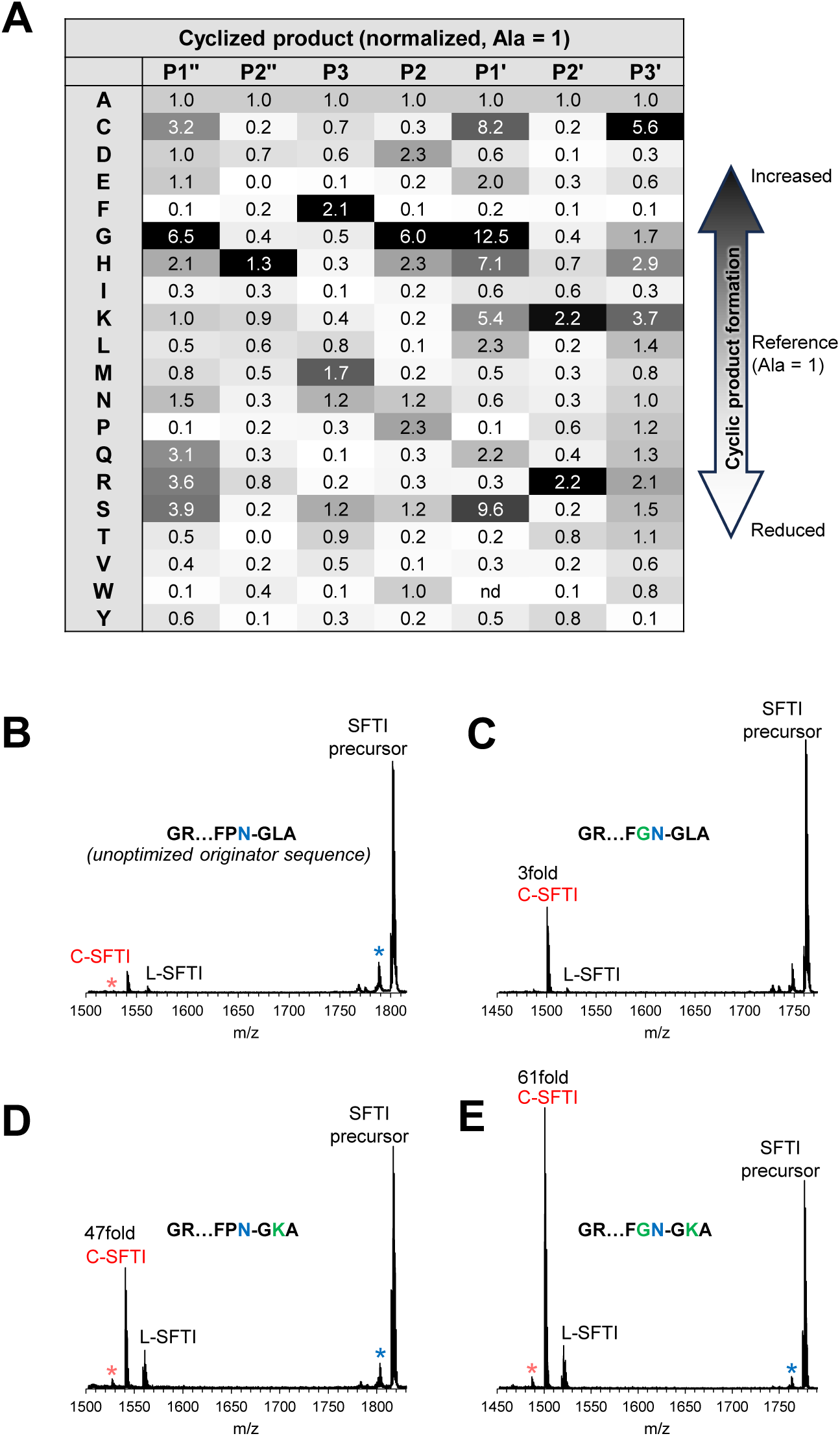
Human legumain prefers positively charged amino acids at position P2’ and glycine at position P1’ and P1’’ for efficient ligation. **(A)** Results of the positional peptide scanning assay using a crude peptide library of 140 variants are summarized in a heatmap. The relative amount of cyclic SFTI product formed was normalized to the peptide containing alanine at the respective position. Dark grey indicates increased cyclic product formation relative to alanine; white indicates decreased formation. **(B-E)** MALDI-ToF mass spectra of selected SFTI variants after incubation with human legumain (hLEG): **(B)** unoptimized G¹R…FPN¹⁴GLA peptide, **(C)** G¹R…FG¹³N¹⁴GLA peptide, **(D)** G¹R…FPN¹⁴GK¹⁶A peptide, and **(E)** G¹R…FG¹³N¹⁴GK¹⁶A peptide. The indicated peptides were resynthesized at >95% purity. Peaks corresponding to the unprocessed SFTI precursor, the cyclic SFTI product (C-SFTI), and the linear intermediate lacking residues G¹⁵LA¹⁷ or G^15^KA^17^ (L-SFTI) are annotated. A blue star indicates a peak corresponding to a demethylated (Δ14 Da) SFTI-GLA species, while a light-red star marks a demethylated (Δ14 Da) c-SFTI species. This modification likely occurs at Thr⁴ or Leu¹⁶ and is frequently observed in our MS measurements ^27^.

Previous studies on plant legumains suggested that proline at P2 impedes cleavage of the cyclic product due to cis-trans isomerization during cyclization ^28^. We found proline to be indeed among the preferred residues for cyclization. However, in our positional scanning assay, glycine at P2 enhanced cyclic product formation threefold compared to proline in hLEG. This suggests that glycine’s increased conformational flexibility may facilitate favorable conformations during the cyclization process, despite the general challenge of cyclic product cleavage.

Collectively, the positional scan revealed that P1’’-Gly, P2’’-His/Lys/Arg/Ala, P3-Phe, P2-Gly/Pro, P1’-Gly, and P2’-Lys/Arg are particularly favorable for hLEG to form the cyclic SFTI product.

### 3. Optimized peptide shows >60fold increase in cyclic product formation

The crude peptide library employed for the initial substrate specificity screen revealed general trends but did not permit quantitative assessment of individual sequence variations. To dissect the contributions of key substitutions and their combinations to cyclization efficiency, we synthesized selected peptide variants at >95% purity and conducted detailed kinetic analyses. The variants tested included P2-P13G (Fig. 2C), P2’-L16K (Fig. 2D), and the double mutant P13G-L16K (Fig. 2E). Each peptide was incubated with hLEG at pH 6.0, and cyclic product formation was quantified by MALDI-TOF MS and compared to the native SFTI-precursor sequence (Fig. 2B). Importantly, all three variants outperformed the native SFTI peptide. The P2-P13G substitution increased cyclic product yield by approximately threefold, suggesting that while Pro at P2 is well tolerated, replacing it with Gly confers a modest benefit. In contrast, the P2’-L16K mutation boosted cyclization by more than 45-fold, underscoring the strong preference for a positively charged residue at the P2’ position. Remarkably, the P13G-L16K double mutant displayed a synergistic effect, producing over 60-fold more cyclic product than the wild type.

To further probe specificity at the P2″ position, we evaluated the substitutions P2″-R2A (Fig. S1A) and P2″-R2H (Fig. S1B), as well as the corresponding triple mutants R2A-P13G-L16K (Fig. S1C) and R2H-P13G-L16K (Fig. S1D). None of these variants showed increased cyclic product formation relative to peptides retaining Arg at P2″, indicating that this position strictly favors arginine/lysine and does not contribute positively to further optimization.

From these results, we derived the optimized sequence motif G^1^-R/K-…-F-G-N^14^-G-R/K-X designated SFTI_opt_, which represents a highly efficient substrate for hLEG-mediated peptide cyclization.

### 4. SFTI_opt_ follows a homologous cooperativity binding mode

Time-course experiments with the optimized substrate SFTI_opt_ revealed that ∼80% of the cyclic product was generated within 5 min of incubation with hLEG (Fig. 3A, S2A). To determine whether this rapid plateau was due to enzyme instability, hLEG was preincubated at either pH 4.0 (optimal for thermal stability) or pH 6.0 (assay buffer pH) for 2 min at 37 °C prior to the reaction (Fig. S2A). Legumain preincubated at pH 6.0 was inactive, indicating that the enzyme remains active for only a short period of time under assay conditions. By contrast, preincubation at pH 4.0 preserved activity, and maximal cyclic product formation was reached within ∼5 min, defining the effective stability window of hLEG during the assay (Fig. S2A).

**Figure 3.**
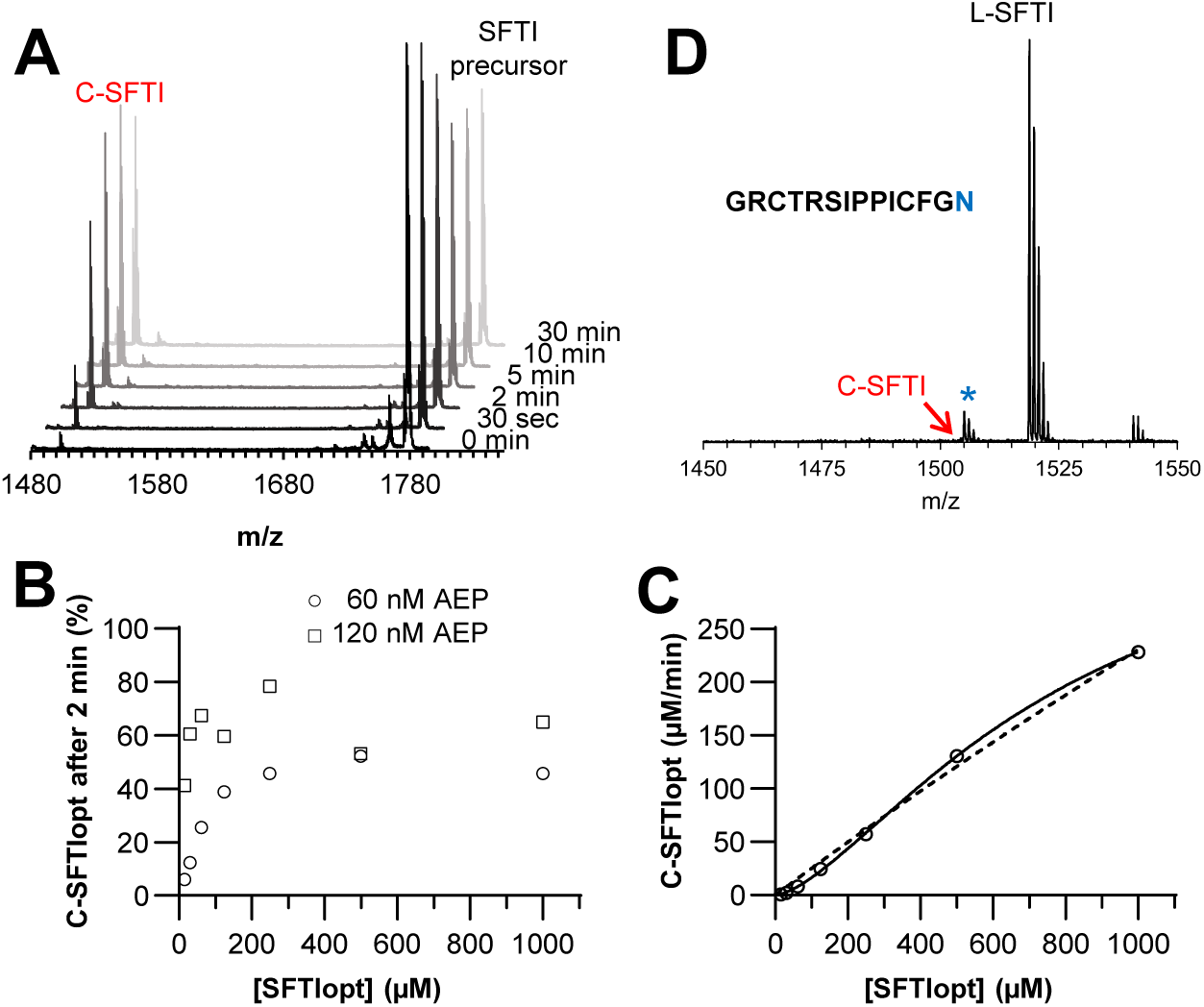
Turnover of SFTI_opt_ is governed by the conformational stability of legumain and does not follow the classical Michaelis-Menten binding mode. **(A)** MALDI-ToF mass spectra of the SFTI_opt_ peptide after incubation with hLEG at the indicated time points. The cyclic reaction product is labeled as C-SFTI. **(B)** Relative amount of cyclized SFTI_opt_ (C-SFTI_opt_), expressed as the percentage of total SFTI in the reaction, at increasing substrate concentrations after 2 min incubation with 60 nM or 120 nM hLEG. **(C)** Kinetic profile of hLEG–catalyzed SFTI cyclization.

Given this stability constraint, we hypothesized that product yield would also be dependent on enzyme concentration. Indeed, incubation of increasing SFTI_opt_ concentrations with either 60 nM or 120 nM hLEG showed that higher enzyme concentrations enhanced cyclic product formation, particularly at low substrate concentrations (Fig. 3B).

To assess the binding characteristics of hLEG toward SFTI_opt_, cyclic product formation was measured over a broad substrate concentration range. The resulting curve did not follow classical Michaelis-Menten kinetics, but instead displayed a sigmoidal shape, with a slow increase at low substrate concentrations (<200 µM), followed by a sharp increase at higher concentrations (Fig. 3C). Fitting the data with an allosteric sigmoidal model yielded a substantially improved fit. This kinetic behavior is consistent with positive cooperativity and suggests the presence of multiple substrate-binding events ^29^. This interpretation was supported by an Eadie-Hofstee transformation, which exhibited the diagnostic curvature associated with homologous cooperativity ^30^ (Fig. S2B). Positive homologous cooperativity refers to a scenario in which binding of one substrate molecule enhances the binding of additional substrate molecules. The molecular basis for such behavior in the case of SFTI is not immediately obvious. However, an alternative explanation is a “memory effect,” in which the catalytic turnover of one SFTI molecule leaves hLEG in a transiently altered state that facilitates subsequent substrate binding. This phenomenon would constitute kinetic cooperativity rather than classical allosteric cooperativity and is consistent with the observed concentration dependence, where the effect becomes more pronounced at higher substrate concentrations. Regardless of the underlying mechanism, these results demonstrate that SFTI cyclization by human legumain does not follow classical Michaelis–Menten kinetics. The dashed line represents the fit of the data to the Michaelis-Menten model, while the solid line represents the fit to the allosteric sigmoidal model. Curve fitting was performed using GraphPad Prism 9.3.1. **(D)** MALDI-ToF mass spectrum of the G¹…N¹⁴ SFTI peptide after incubation with hLEG. No cyclic product formation was detected.

### 5. SFTI_opt_ is a substrate for transpeptidase activity but not for ligase activity

Our previous work showed that fully activated hLEG converts the SFTI precursor to the cyclic product predominantly via transpeptidation, with little or no conversion of the linear product (L-SFTI) to the cyclic form by ligation. While ligase activity has been reported for hLEG on certain substrates, such as synthetic linear peptides and cystatin E ^22, 23^, this reaction appears to be highly substrate-dependent. The original SFTI precursor sequence may therefore have been suboptimal for ligation. Having developed the optimized SFTI_opt_ sequence, we next tested whether it could serve as a ligase substrate. For this, we used the linear L-SFTI_opt_ peptide, which retains Gly at P2 but lacks the prime-side Gly^15^–Lys^16^–Ala^17^ residues present in the precursor. When incubated with hLEG, this linear peptide showed no detectable cyclization (Fig. 3D). These results confirm that, even in its optimized form, the SFTI scaffold is not well-suited to hLEG-mediated ligation, and that cyclic product formation in this system arises primarily through transpeptidase activity.

### 6. Interaction of P2’/P2’’ residue with Asp160 is critical for efficient transpeptidase activity

To explore why specific sequence variations enhanced cyclization of the SFTI precursor by hLEG, we used AlphaFold 3 to model hLEG-peptide complexes ^31^. The complex of hLEG with the full-length SFTI precursor could not be reliably modeled as judged by the positioning of the P1-Asn in the S1 pocket; therefore, we focused on short substrate-mimetic peptides comprising only positions P3–P2–P1–P1’–P2’. The sequences modeled were FPNGL, FGNGK, FGNGR, FPNGK, and FPNGR (Fig. 4 and S3A,B). The models revealed a distinct difference in binding at the prime-side substrate site: peptides with hydrophobic Leu at P2’ adopted a different orientation compared to those with positively charged Lys or Arg. In the latter, the P2’ side chain formed favorable electrostatic interactions with Asp160 in the S2’ pocket of hLEG.

**Figure 4.**
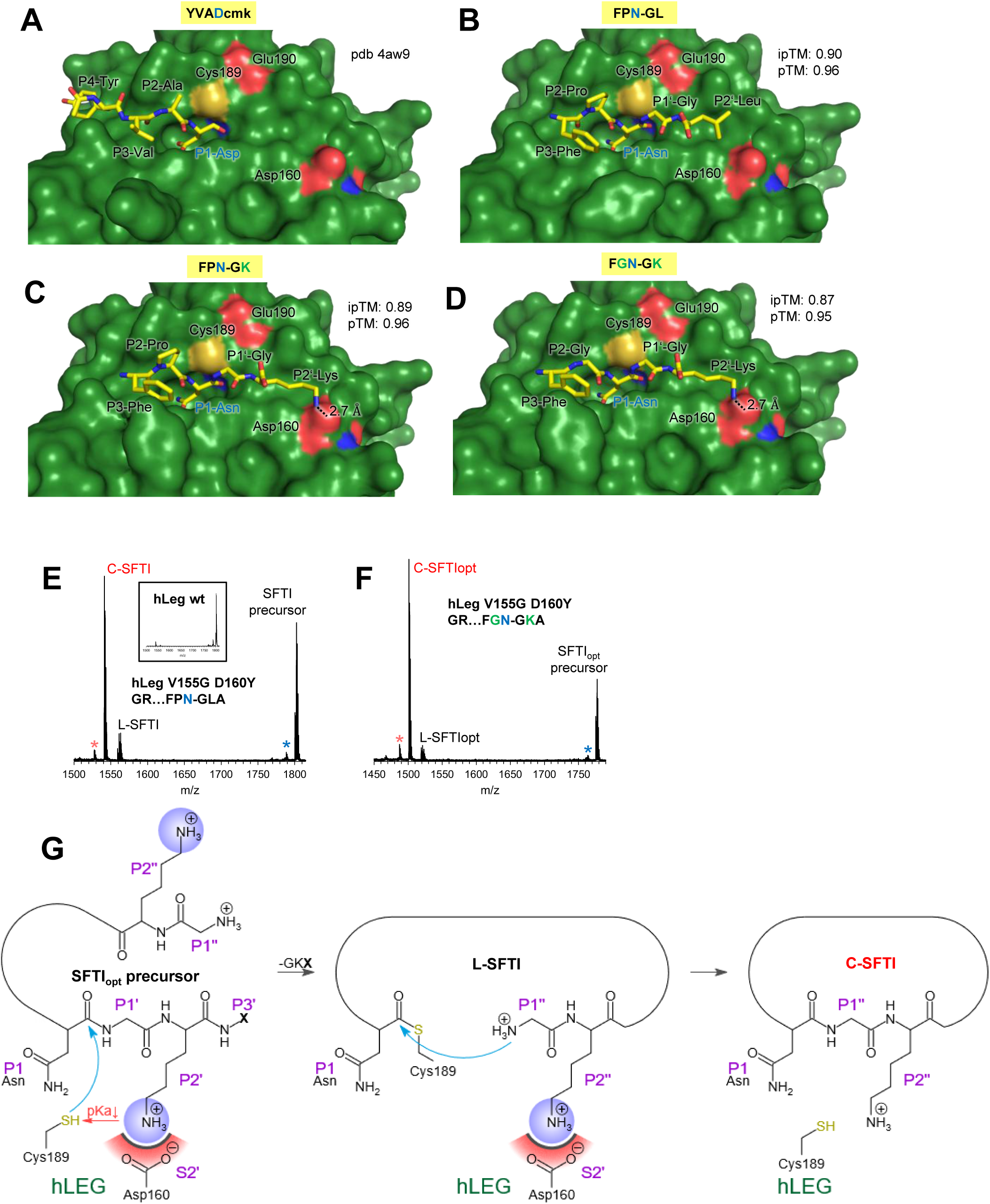
Asp160 in the S2′ pocket of human legumain is a key determinant of substrate specificity. **(A)** Crystal structure of hLEG in complex with the covalent inhibitor YVAD-cmk (Tyr-Val-Ala-Asp–chloromethylketone) (PDB: 4AW9). **(B–D)** AlphaFold 3 models of hLEG bound to the indicated peptides: **(B)** FPN¹⁴–GL, **(C)** FPN¹⁴–GK, and **(D)** FGN¹⁴–GK. **(E)** MALDI-ToF mass spectrum of the unoptimized G¹…N¹⁴–GLA SFTI precursor peptide after incubation with the V155G–D160Y mutant of hLEG. The corresponding spectrum for the reaction catalyzed by wild-type hLEG is shown in the inset. **(F)** MALDI-ToF mass spectrum of the SFTI_opt_ precursor peptide after incubation with the V155G–D160Y variant of hLEG. A blue star indicates a peak corresponding to a demethylated (Δ14 Da) SFTI–GLA species, and a red star marks a demethylated (Δ14 Da) c-SFTI species. **(G)** Reaction scheme of the SFTI-cyclisation by hLEG. Protonation of Cys189 is regulated by the P2’ residue. Positively charged P2′/P2′′ residues may facilitate deprotonation of Cys189, either directly or indirectly by compensating the negative electrostatic influence of the nearby Glu190. In step 1 of the transpeptidation reaction the Cys189 Sγ is attacking the scissile peptide bond leading to the formation of a thioester intermediate. In Step 2 of the reaction the intermediate is released by the N-terminal P1’’ residue.

To test the functional relevance of this interaction, we examined the V155G-D160Y mutant of hLEG, engineered to mimic the S2′ pocket of *Arabidopsis thaliana* legumain (AtLEG). Using the unmodified SFTI precursor as substrate, this mutant displayed higher turnover and greater cyclic product yield than wild-type hLEG (50 % vs. 10 %, Fig. 4E). Substituting the precursor with SFTI_opt_ further increased the proportion of cyclic product by approximately 20 % (from 53 % to 73 %) (Fig. 4F). By contrast, wild-type hLEG produced an approximately 60-fold increase in cyclic product yield when catalyzing cyclization of SFTI_opt_ relative to the unoptimized precursor (Fig. 2E).

The modest enhancement observed for the V155G-D160Y mutant, compared to the much larger effect in wild-type hLEG, suggests that SFTI_opt_ was specifically tuned to the native S2′ pocket of hLEG, rather than an AtLEG-like configuration. Nevertheless, the fact that the ligase activity still increased, contrary to expectations based solely on binding properties, indicates that an Arg residue at P2′ exerts a positive catalytic effect, likely through *<u>k</u>*_cat_ modulation. Mechanistically, a positively charged P2′ residue appears to enhance catalytic efficiency by facilitating deprotonation of the catalytic cysteine (Cys189), either by lowering its p*K*a directly or by counteracting the negative electrostatic effect of the nearby Glu190. This, in turn, increases nucleophilicity and promotes formation of the thioester intermediate (Fig. 4G). Consistent with these experimental observations, AlphaFold 3 models of the hLEG-V155G-D160Y mutant bound to FGNGL or FGNGK peptides revealed that the FGNGL complex achieved a high ipTM score (0.9), whereas the FGNGK complex scored poorly (0.49) and misplaced the P2′-Lys into the S1 pocket instead of correctly positioning the P1-Asn (Fig. S3C,D).

Collectively, these results demonstrate that interactions between the P2′/P2′′ residues and Asp160 are key determinants of hLEG substrate specificity and transpeptidase activity.

### 7. AtLEGβ prefers Leu in both P2’ and P2”

To further assess the contribution of the S2’ pocket to substrate recognition, we compared the activity of *Arabidopsis thaliana* legumain isoform β (AtLEGβ) with hLEG (hLEG) (Fig. 5). AtLEGβ possesses an S2’ pocket optimized to accommodate hydrophobic Leu/Ile side chains. Using the wild-type SFTI precursor peptide as a substrate, AtLEGβ exhibited a higher turnover rate and approximately threefold higher yield of cyclic product compared to hLEG at equal enzyme concentrations (Fig. 5A). In detailed positional scanning experiments, AtLEGβ showed highest turnover toward cyclic product formation when Leu was present at both P2’ and P2’’, resulting in almost 100% conversion to the cyclic product (Fig. 5B). Only ∼40% conversion was observed when Arg occupied the P2’’ position, which corresponds to the unmodified SFTI sequence. When Arg was present at both P2’ and P2’’, turnover improved compared to Arg only at P2’’ (Fig. 5C), suggesting that also here the P2’ residue facilitates deprotonation of the catalytic Cys211, thereby promoting thioester formation. Although the S2’ pocket of AtLEGβ is optimized for hydrophobic residues, the enzyme tolerated Arg at P2’, supporting a *k*_cat_-tuning effect at this position. In contrast, hLEG showed highest activity when Arg was present at both P2’ and P2’’, consistent with both optimal electrostatic complementarity in the S2’ pocket and *k*_cat_ tuning. Substitution of Arg to Glu at P2’’ resulted in almost 100% conversion to the cyclic product by AtLEGβ, likely due to favorable electrostatic interaction of P2’’ Glu with His182 in the S2’ pocket of AtLEGβ (Fig. 5D).

**Figure 5.**
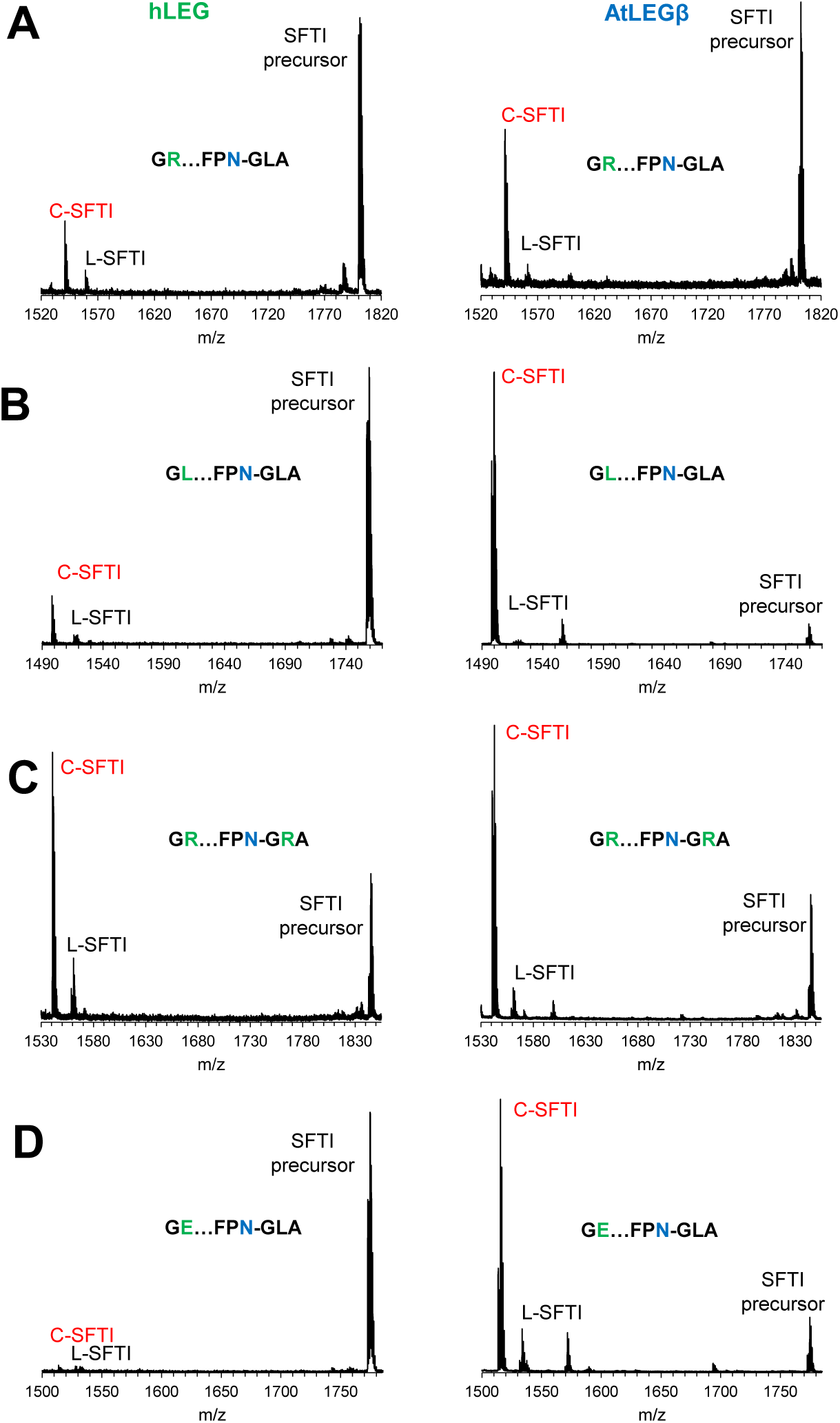
**AtLEGβ prefers leucine both in position P2’ and P2’’**. **(A-D)** MALDI-ToF mass spectra of selected SFTI variants after incubation with human legumain (hLEG) or AtLEGβ. **(A)** unoptimized G¹R…FPN¹⁴GLA peptide, **(B)** G¹L…FG¹³N¹⁴GLA peptide, **(C)** G¹R…FG¹³N¹⁴GLA peptide, and **(D)** G¹E…FG¹³N¹⁴GLA peptide. Peaks corresponding to the unprocessed SFTI precursor, the cyclic SFTI product (C-SFTI), and the linear intermediate lacking residues G¹⁵LA¹⁷ (L-SFTI) are annotated.

These findings highlight distinct specificity determinants in the S2’ and S2’’ subsites of plant and human legumain and suggest that positional scanning could be further exploited to optimize SFTI-based substrates for other legumains.

### 8. Legumain’s total catalytic activity shows similar sequence preferences on the SFTI scaffold

Because the SFTI precursor can also undergo hydrolysis by hLEG, we used our positional scanning dataset to examine sequence preferences in this reaction mode. Both linear L-SFTI and cyclic C-SFTI formation proceed through a common thioester intermediate: either by attack from a water molecule (leading to hydrolysis) or from the peptide’s N-terminus (resulting in cyclization, aminolysis). Accordingly, total lytic activity was quantified based on the combined yield of linear and cyclic products (Fig. 6). When total product formation was analyzed, substrate preferences for peptide lysis closely matched those for transpeptidation: Phe at P3, Gly or Pro at P2, Gly at P1’, and basic residues (Lys, Arg) at P2’. P3’ showed low specificity. Accordingly, the optimized sequence for hydrolysis under this criterion was identical to that for transpeptidation. This is consistent with the fact that the attack by the catalytic Cys189 residue is the rate-limiting step in both hydrolysis and transpeptidation reactions.

**Figure 6.**
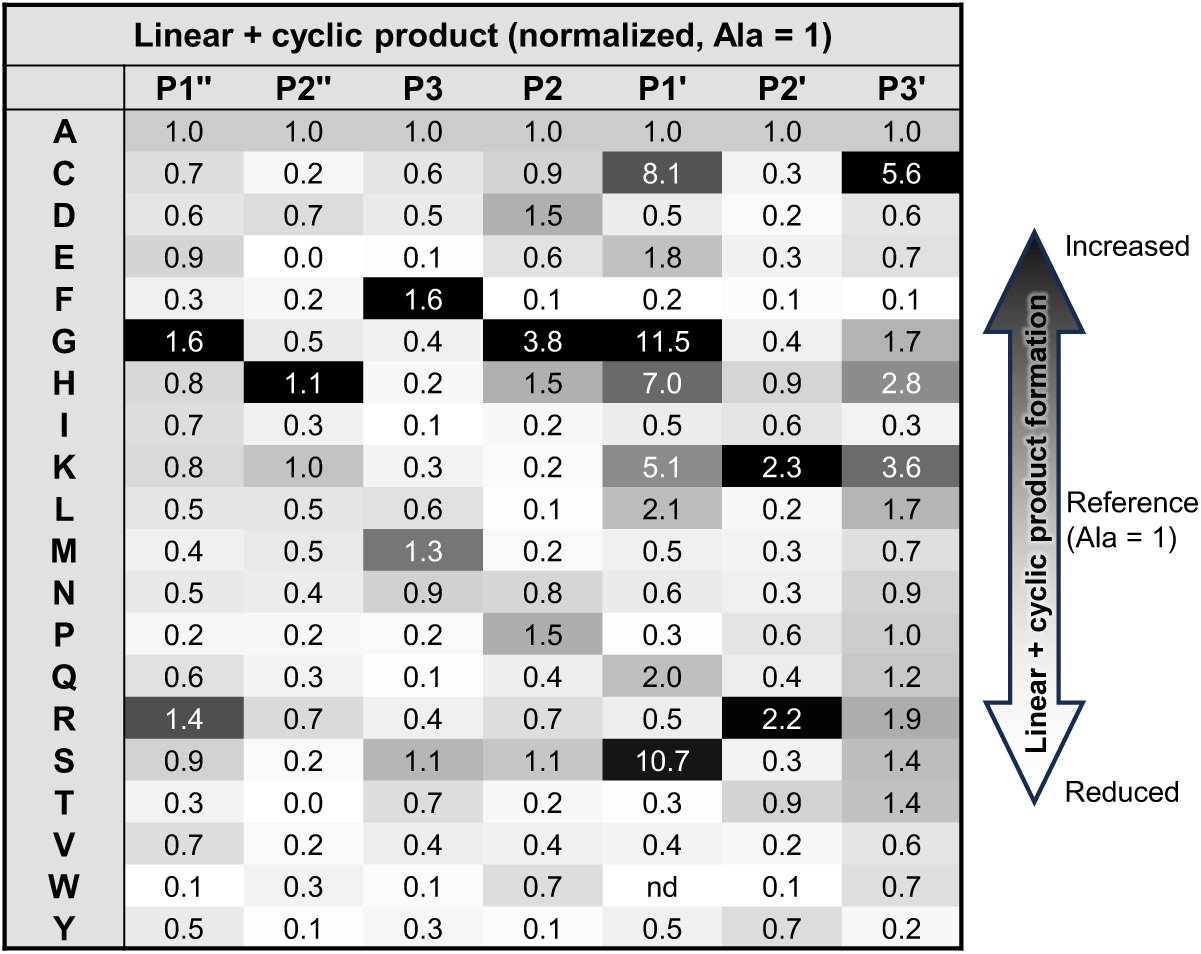
Legumain’s total lytic activity exhibits similar sequence preferences on the SFTI scaffold. Results of the positional peptide scanning assay using a crude peptide library of 140 variants are summarized in a heatmap. The relative amount of total SFTI product (cyclic + linear) formed was normalized to the peptide containing alanine at the respective position.

Interestingly, when the analysis was restricted to linear product formation, *i.e.* hydrolysis exclusively, distinct preferences emerged, especially at P1’, P2, and P3 (Fig. S4). While Gly was favorable for total product formation (cyclic + linear), Ser was preferred at P1’ when only hydrolysis was considered. At position P2, residues such as Arg, Cys, Glu, Ala, and Ser were favored in the context of linear product formation, in contrast to the preference for Gly when cyclic + linear products were analyzed. Notably, glycine was even unfavourable at P2 position when the formation of the linear product was analyzed. P3 also showed broader specificity in hydrolysis, with Ala, Arg, Ser, and Val being tolerated, whereas Phe was clearly favored when analyzing total product formation.

These findings may suggest that substrates which are poor transpeptidation candidates tend to be better hydrolysis substrates. However, this interpretation is likely misleading, as the catalytic Cys189 attack remains the rate-limiting step in both processes. Steric properties of the substrate likely determine whether the intermediate is resolved via intramolecular transpeptidation or hydrolysis by a water molecule. In any way, for a good transpeptidation substrate a relatively long residence time of the cleaved, non-primed product is important to give the incoming new amino terminus (P1’’) enough time for productive aminolysis. By contrast, residence time should be minimal for an exclusive hydrolysis substrate.

### 9. *In silico* specificity profiling reproduced experimental specificity results

The SFTI precursor imposes steric constraints that are likely to influence hLEG’s substrate preferences, meaning that specificity for linear peptides may differ from that observed with the SFTI scaffold. To gain further insight, we performed an *in silico* substrate profiling using AlphaFold 3. We modeled active hLEG in complex with a series of hexapeptides spanning both non-prime and prime substrate-binding sites, thereby probing specificity in the protease and transpeptidase mode. The initial design was based on the YVAN–XX (P4 – P2’) substrate sequence, chosen to evaluate prime-side preferences. YVAN was selected for the non-prime segment because crystallographic studies have shown it binds favorably to hLEG. In particular, the structure of hLEG bound to the inhibitor YVAD-cmk (PDB: 4AW9) revealed stabilizing interactions with the non-prime substrate binding sites. To systematically explore prime-side preferences, we substituted the two C-terminal residues (XX) in YVAN-XX with all 20 proteinogenic amino acids, thereby generating 400 distinct peptide-hLEG complexes. Each model was evaluated using the inter-protein Template Modelling-score (ipTM), which measures the predicted accuracy of protein–peptide orientation. Scores >0.8 indicate high-confidence predictions, whereas values <0.6 reflect low confidence or likely modeling failure ^31^. We used the ipTM score as a proxy to estimate the affinities of the AF3-predicted ligand enzyme complexes. Most complexes scored above 0.8, suggesting broad steric tolerance at P1′ and P2′ (Fig. 7A). However, consistent trends emerged. Positively charged residues such as Lys and Arg were strongly favored at P2′, mirroring the preferences observed experimentally in our SFTI-based positional scanning assay. Overall, Pro was the most favored residue at P2’ *in silico* for linear substrates, but Tyr was also well tolerated. Arginine or lysine at P2′ may enhance catalytic efficiency by facilitating deprotonation of the catalytic cysteine (Cys189), either directly or indirectly through interactions with the nearby Glu190, which otherwise destabilizes the thiolate form of Cys189 by increasing its p*K*a. The presence of a positively charged residue could thus lower the effective p*K*a of Cys189, enhancing its nucleophilicity and promoting thioester intermediate formation. In this way, electrostatic compensation by P2′ residues may counteract the inhibitory influence of Glu190 on catalysis.

**Figure 7.**
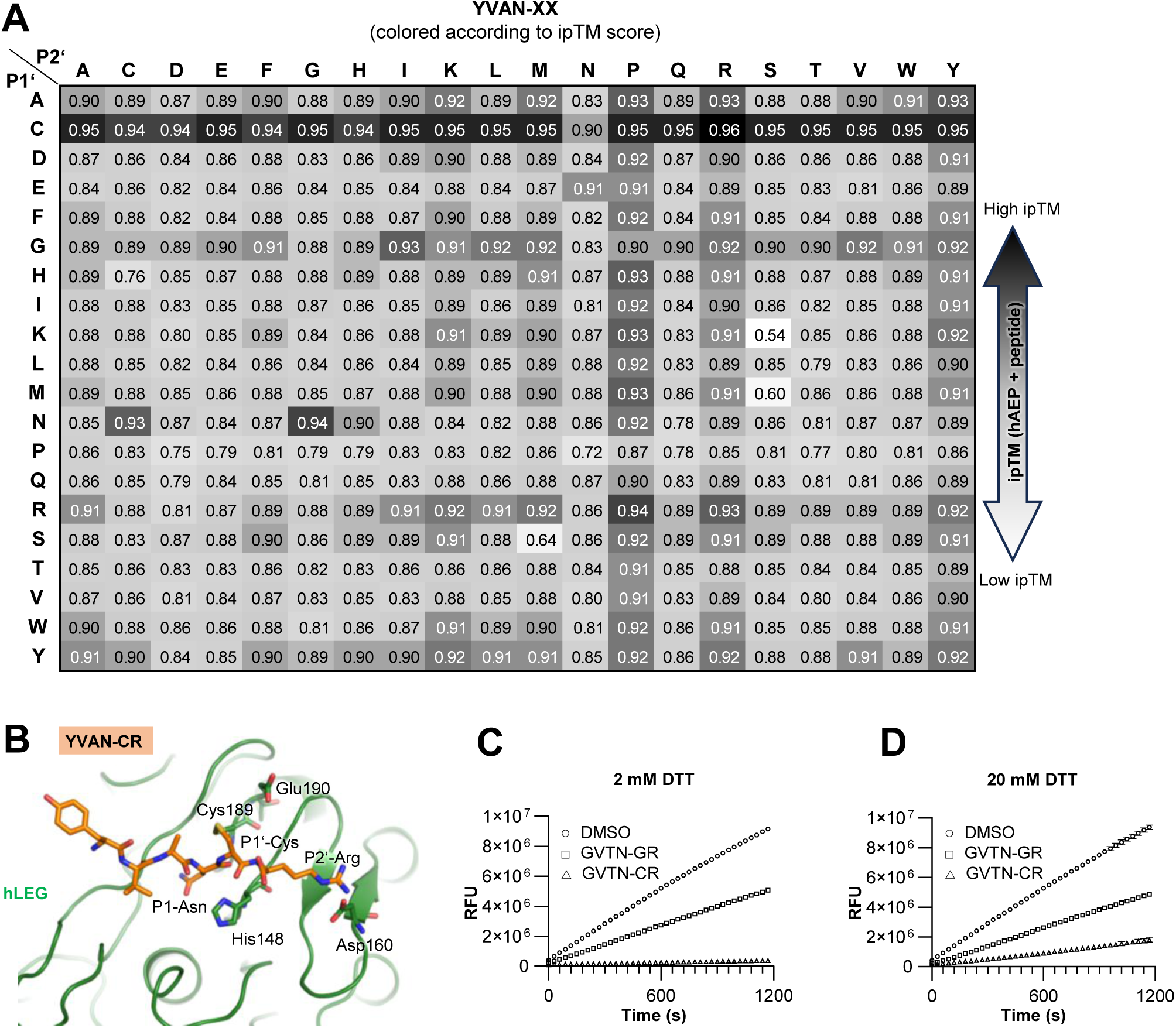
*In silico* substrate specificity screening of the prime side of human legumain reproduces its preference for basic residues at P2′ and identifies P1′ cysteine as a covalent inhibitor. (**A**) Results of the AlphaFold 3-based *in silico* substrate specificity screening using the sequence of hLEG and the YVAN-XX peptide library, summarized in a heatmap colored according to the ipTM score of the respective complexes. **(B)** AlphaFold 3 model of hLEG bound to the YVAN-CR peptide, which achieved the highest ipTM score (0.96) in the *in silico* screen. The disulfide bond formed between the P1′ cysteine of the peptide and the catalytic Cys189 of hLEG is shown. **(C, D)** Competition assays of hLEG with the peptides GVTN-CR and GVTN-GR. Each peptide (2 mM) was preincubated with the fluorogenic substrate Z-AAN-AMC in assay buffer (pH 5.5) containing either 2 mM DTT **(C)** or 20 mM DTT **(D)**. Reactions were initiated by the addition of 2 nM hLEG, and fluorescence was recorded over 20 min at 37 °C.

At P1′, Gly was preferred, also consistent with our experimental findings, though Cys produced models with even higher ipTM scores. Closer inspection of the P1′-Cys models revealed a predicted disulfide bond between the P1′ cysteine and hLEG’s catalytic Cys189 (Fig. 7B). This observation aligns with previous reports of covalent disulfide formation between Cys189 and P1′ cysteine in macrocypin inhibitors and short peptide substrates ^23, 32^.

### 10. Cysteine in P1’ functions as a covalent hLEG inhibitor

Our *in silico* specificity screen of hLEG’s prime-side binding site suggested that peptides with cysteine at P1′ could act as covalent inhibitors by forming a disulfide bond with Cys189, the enzyme’s catalytic residue. To test this, we performed a competition assay in which hLEG was incubated with the fluorogenic substrate Z-AAN-AMC (25 µM) in the presence of the GVTN-CR peptide (2 mM) (Fig. 7C). Controls included DMSO or the GVTN-GR peptide, which lacks the P1′ cysteine. In the presence of GVTN-CR, substrate turnover was completely inhibited, consistent with covalent active-site binding. By contrast, GVTN-GR reduced activity by only ∼50%, likely through non-covalent competition at the substrate binding site. To determine whether this inhibition was disulfide-dependent, we repeated the assay under reducing conditions (20 mM DTT) (Fig. 7D). Under these conditions, GVTN-GR inhibition remained unchanged, whereas GVTN-CR inhibition was markedly reduced, confirming that the covalent interaction is mediated by disulfide bond formation between the P1′ cysteine and Cys189 on hLEG (Fig. S5).

Interestingly, both the SFTI-based screening and the *in silico* analysis independently highlighted cysteine at P1′ as functionally significant. In the SFTI-based assay, P1′-Cys supported efficient cyclic product formation, and *in silico* modeling predicted a potential covalent interaction between P1′-Cys and Cys189. However, during SFTI cyclization, direct disulfide bond formation between these residues is unlikely. Although a disulfide could theoretically form prior to nucleophilic attack, this scenario is unlikely because the active site is optimized for catalyzing nucleophilic reactions rather than promoting disulfide exchange. Thus, the nucleophilic attack is expected to proceed preferentially. Alternatively, the P1′ cysteine may simply form favorable non-covalent interactions within the S1′ pocket without disulfide formation. This interpretation is supported by the observation that P1′-Ser performs comparably to P1′-Cys in promoting cyclic product formation. These findings suggest that P1′-Cys may serve as a covalent “warhead”, providing a starting point for the design of covalent peptide-based inhibitors. This concept could be extended to other cysteine proteases, including caspases, metacaspases, and paracaspases. Notably, we also performed the AlphaFold-based screen for human caspase-9, where however a Cys at P1′ was not predicted to form a disulfide bond, likely due to steric constraints characteristic of the caspase S1′ pocket ^33^.

### 11. Substrate specificity is broader on the non-prime side than on the prime-side

To probe hLEG’s preferences at upstream (non-prime) positions, we performed an additional *in silico* substrate screen using peptides of the format XXXN-CR, fixing P1′ as cysteine and P2′ as arginine based on the top-scoring sequences from the prime-side analysis. Specificity trends were visualized in 2D heatmaps, fixing either P4 as tyrosine (Fig. 8A) or P2 as alanine (Fig. 8B). The screen revealed a relatively broad preference at P2 for alanine, cysteine, aspartic acid, serine, and threonine, and at P3 for alanine, glutamic acid, methionine, proline, asparagine, and valine. Although Gly at P2 was strongly favored in the SFTI-based experimental screen, it did not emerge as a top hit here. At P3, ipTM scores were consistently high (>0.90), indicating low selectivity. This trend persisted when P2 was fixed as Ala and P3/P4 were varied (Fig. 8B). The main exception was Cys at P3, which repeatedly produced lower scores across both the YXXN-CR and XXAN-CR screens. For P4, Gly and Val were associated with the highest-scoring models (ipTM up to 0.97), although most P4 substitutions yielded scores >0.90, suggesting broad tolerance. The overall pattern of permissiveness at P3 and P4 aligns well with results from an experimental combinatorial fluorogenic substrate library screen ^34^.

**Figure 8.**
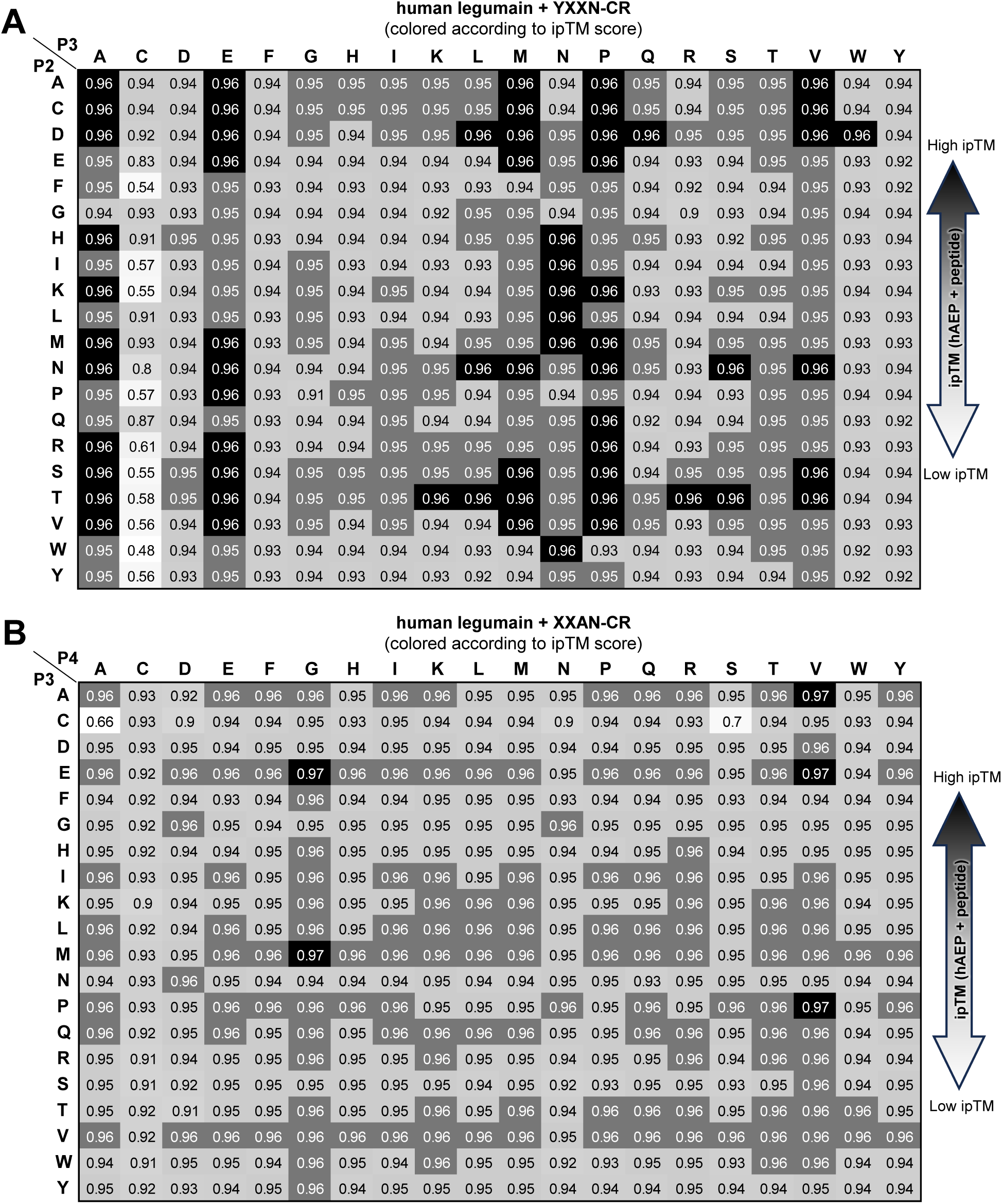
Substrate specificity is relatively broad on the non-prime side of human legumain. Results of the AlphaFold 3-based *in silico* substrate specificity screening using the sequence of hLEG and the XXXN-CR peptide are shown. To enable two-dimensional visualization, either the P2 position was fixed as alanine **(A)** or the P4 position was fixed as tyrosine **(B)**. Heatmaps are colored according to the ipTM score of the respective legumain–substrate complexes.

Overall, ipTM scores tended to be higher on the non-prime side, implying greater sequence flexibility in this region compared to the prime side. The main constraints were the requirement for Asn or Asp at P1 and the poor compatibility of Cys at P3. From these results, we derived an optimized substrate/inhibitor motif for hLEG: X-P/A/V/E-A/T/D/S-N-C/G-R/K. Given that our SFTI-based positional scanning identified similar specificity patterns for both transpeptidase and hydrolase activity, we conclude that the preferences identified here for linear peptides can be extrapolated to hLEG’s hydrolase as well as its transpeptidase activity.

### 12. *In silico* substrate specificity-screening enables the development of optimized hLEG substrates

To evaluate whether *in silico* substrate specificity profiling can guide the design of improved hLEG substrates, we selected tripeptide-AMC fluorogenic substrates for experimental validation. We compared Z-AAN-AMC, a well-established reporter of hLEG protease activity, with Z-VAN-AMC, a less commonly used analogue. Although ipTM scores for Ala and Val at P3 were similarly high, experimental substrate turnover assays revealed notable differences. Z-AAN-AMC exhibited a K_m_ of 55 µM, reflecting moderate affinity for hLEG (Fig. 9A). In contrast, Z-VAN-AMC showed a K_m_ of ∼20 µM, indicating stronger binding, and displayed substrate inhibition with a K_i_ of 16 µM (Fig. 9B). Most strikingly, V_max_ for Z-VAN-AMC was ∼20-fold higher than for Z-AAN-AMC (7800 vs. 400 RFU/s), underscoring its markedly superior catalytic efficiency.

**Figure 9.**
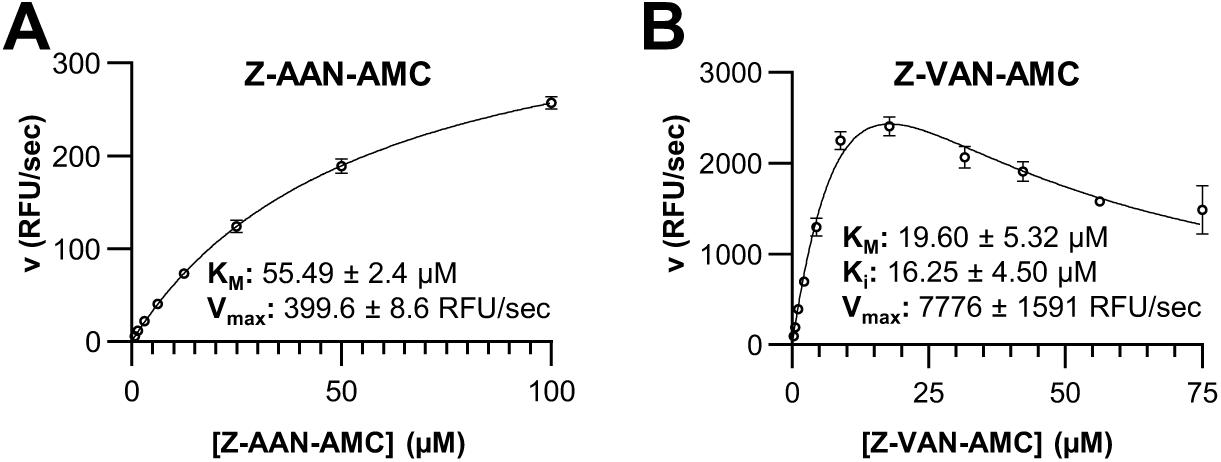
Kinetic characterization of Z-AAN-AMC and Z-VAN-AMC substrates reveals enhanced turnover of Z-VAN-AMC, consistent with *in silico* substrate specificity predictions. (**A**) Kinetic profile of hLEG-catalyzed Z-AAN-AMC cleavage. The solid line represents the fit of the data to the Michaelis–Menten model. **(B)** Kinetic profile of hLEG-catalyzed Z-VAN-AMC cleavage. The solid line represents the fit of the data to the substrate inhibition model in GraphPad Prism 9.3.1.

13. *In silico* substrate specificity screening is transferable across legumain family proteases To evaluate whether our *in silico* substrate profiling strategy can be applied to other proteases, we tested it on AtLEGβ. Following the same workflow used for hLEG, we generated a peptide library of the format AAAN-XX, fixing Asn at P1 (essential for catalysis) and systematically varying P1′ and P2′ across all 20 proteinogenic amino acids. Strikingly, AtLEGβ displayed a strong preference for cysteine at P1′, mirroring our findings for hLEG (Fig. 10A). In these models, P1′ cysteine formed a predicted disulfide bond with the catalytic Cys211 (Fig. 10B), consistent with the covalent inhibition mechanism proposed for hLEG. To test whether this disulfide-mediated inhibition mechanism also operates in AtLEGβ, we performed competition assays using the GVTN-GR and GVTN-CR peptides. Notably, GVTN-CR caused complete inhibition of AtLEGβ activity (Fig. 10C), which was partially reversed upon addition of 20 mM DTT (Fig. 10D and Fig. S6), confirming a covalent, disulfide-mediated inhibitory mode.

**Figure 10.**
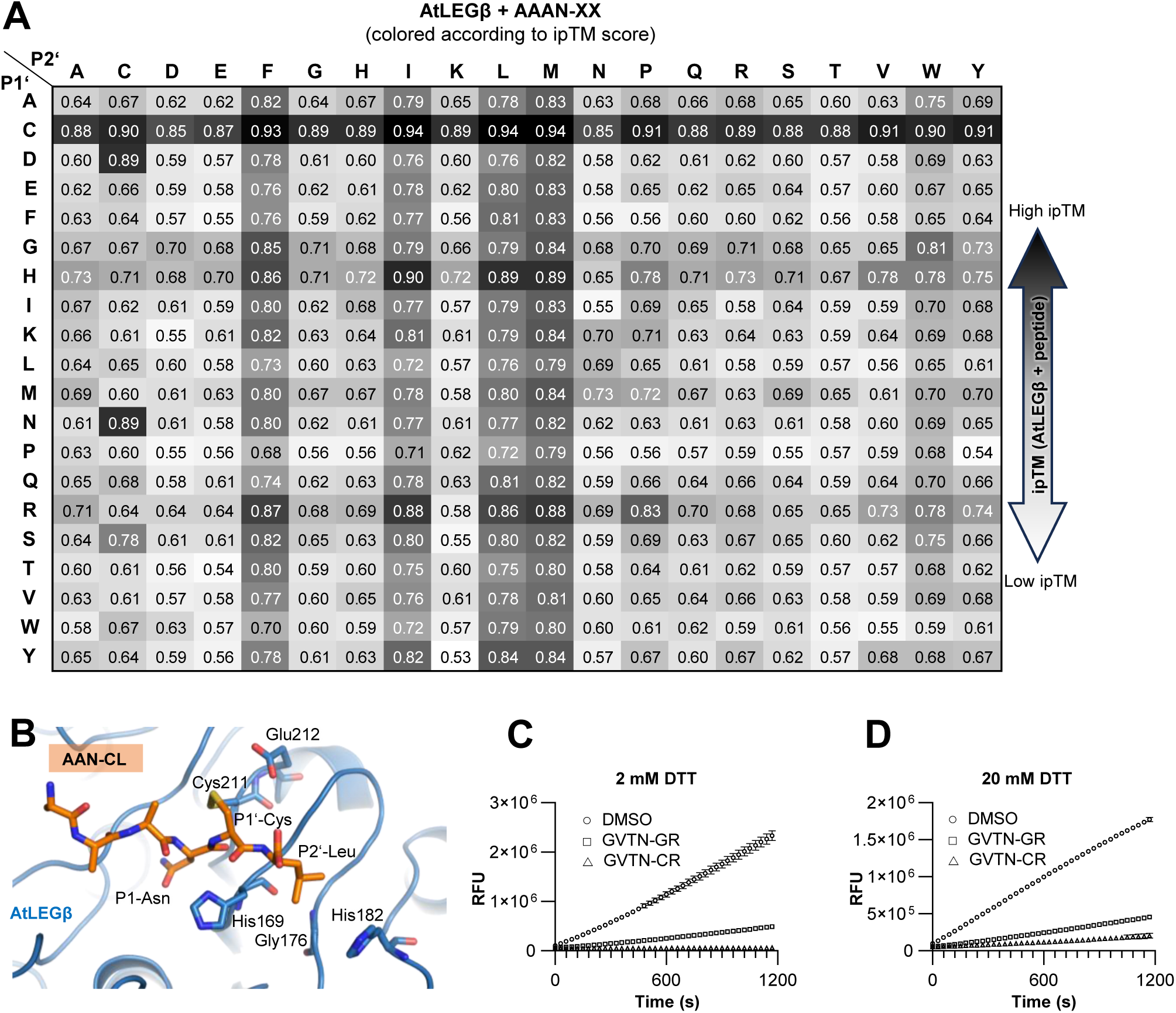
*In silico* substrate specificity screening of the prime side of *Arabidopsis thaliana* legumain β (AtLEGβ) reveals a preference for hydrophobic residues at P2′ and identifies P1′ cysteine as a covalent inhibitor. (**A**) AlphaFold 3-based *in silico* substrate specificity screening of AtLEGβ using the AAAN-XX peptide library, summarized in a heatmap colored by ipTM score. **(B)** AlphaFold 3 model of AtLEGβ bound to the top-ranking AAAN-CL peptide (ipTM 0.94), showing a disulfide bond between the P1′ cysteine and the catalytic Cys211. **(C, D)** Competition assays of AtLEGβ with GVTN-CR and GVTN-GR peptides (2 mM each) using the fluorogenic substrate Z-AAN-AMC in assay buffer (pH 5.5) containing 2 mM DTT **(C)** or 20 mM DTT **(D)**. Reactions were started with 20 nM AtLEGβ, and fluorescence was recorded over 20 min at 37 °C.

At P2′, AtLEGβ showed a clear preference for hydrophobic residues Leu, Ile, Phe, and Met, in excellent agreement with its hydrophobic prime-side pocket and prior PICS-based experimental data ^35^.

To probe non-prime side specificity, we constructed a second peptide library using the format XXXN-CL, fixing P1′ as cysteine and P2′ as leucine, based on the highest-scoring sequences from the prime-side screen. This analysis revealed broad amino acid tolerance at P2 and P3, with notable preferences for Ala, Glu, His, Ile, Thr, and Val at P3 (Fig. S7A). Thr and Val were especially well tolerated, a trend that persisted when P2 was fixed as Ala and P3/P4 were varied (Fig. S7B). These predictions were validated experimentally by comparing AtLEGβ affinity for Z-AAN-AMC versus Z-VAN-AMC fluorogenic substrates. Consistent with the *in silico* results, AtLEGβ showed a significantly lower K_m_ for Z-VAN–AMC than for Z-AAN-AMC (Z-VAN-AMC: 57 ± 3 µM; Z-AAN-AMC: 337 ± 3 µM) ^35^. As with hLEG, cysteine at P3 consistently yielded low ipTM scores, indicating poor compatibility, while most other P3/P4 variants scored above 0.90, reflecting broad tolerance. No clear preference emerged at P4, suggesting high flexibility in this position.

At position P3, the *in silico* screen revealed a strong preference for threonine and valine in AtLEGβ, while hLEG exhibited broader substrate tolerance. In contrast, the SFTI-based screen identified phenylalanine as preferred at P3 in hLEG, an observation not captured in the *in silico* screen. These discrepancies may arise from structural differences between the conformationally constrained SFTI precursors used in the empirical assay and the linear peptides modeled computationally. Such constraints can significantly influence substrate-enzyme interactions and highlight the importance of using context-specific peptide scaffolds in substrate profiling. Both hLEG and AtLEGs possess an Asp147 residue (hLEG numbering) near the active site, which participates in oxyanion hole formation and undergoes conversion to a succinimide (Suc147) in the active enzyme ^23^. However, AlphaFold3 currently lacks the capability to incorporate succinimide residues within polypeptide chains. As a result, our models retained Asp at this position, introducing minor steric clashes in the predicted structures. Despite this limitation, the active sites of both hLEG and AtLEGβ were modeled with high accuracy, yielding overall Cα RMSD values of 0.28 Å for AtLEGβ + AAANCL compared to PDB entry 6YSA, and 0.22 Å for hLEG + YVANCR compared to PDB entry 4AW9. Modeling of AtLEGγ, however, was unsuccessful, highlighting the current challenges in accurately representing post-translational modifications in *in silico* enzyme modeling.

## CONCLUSION

In this study, we employed an SFTI-based substrate specificity screen to probe the transpeptidase activity of hLEG and developed an optimized cyclization substrate SFTI_opt_ with substantially enhanced activity (Fig. 11A). In parallel, we applied an AlphaFold-based *in silico* screening approach as a complementary strategy to identifying improved substrates and study catalytic properties. Both experimental and computational results were highly consistent, underscoring the reliability of combining empirical and structural modeling approaches for mechanistic studies of legumain function.

**Figure 11.**
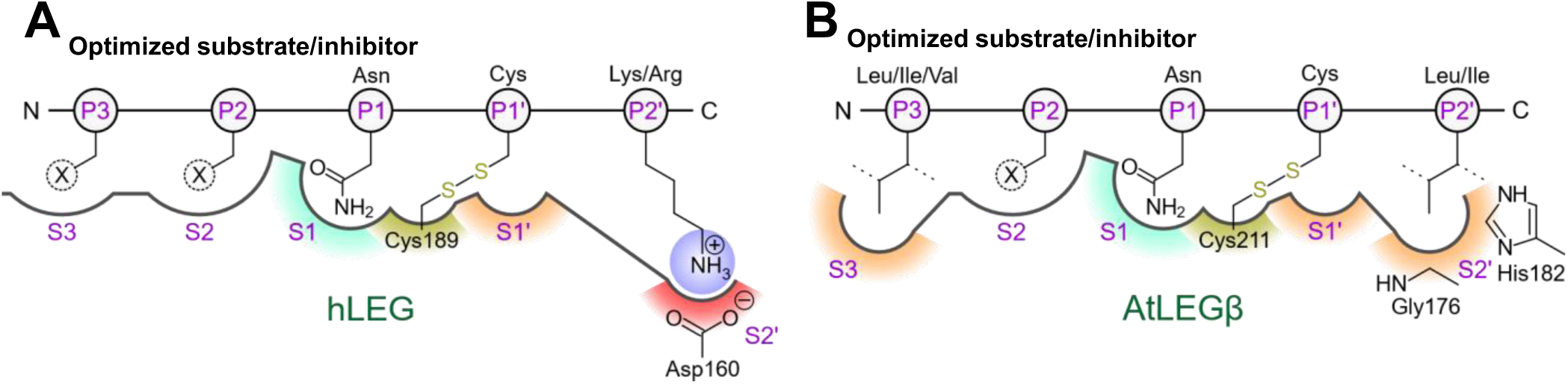
hLEG and AtLEGβ display distinct substrate specificities. **(A)** Human legumain (hLEG) preferentially cleaves/transpeptidates substrates containing basic residues at the P2′ position, while exhibiting a relatively broad substrate tolerance at non-primed positions. At P2, X can be substituted by various amino acids, particularly Ala, Cys, Asp, Ser, and Thr. At P3, substitution of X is most favorable with Ala, Glu, Met, Pro, Asn, and Val. Substrates with a cysteine at P1′ can form a covalent disulfide bond with Cys189 of hLEG, thereby acting as inhibitors. **(B)** AtLEGβ shows a pronounced preference for leucine or isoleucine at the P2′ position and favors small, hydrophobic residues at P3 (Ile, Leu, Val, Ala, Glu, His, Thr). The P2 position is highly flexible. Similarly, substrates containing a cysteine at P1 may become covalent inhibitors through disulfide bond formation with the catalytic Cys211.

Our *in silico* analysis demonstrated strong predictive power for substrate binding and provided valuable mechanistic insights into catalysis. For example, AlphaFold accurately predicted the preference for valine at P3 in AtLEGβ, previously confirmed experimentally (Fig. 11B) ^23^, and helped rationalize differential substrate turnover in human and plant legumains. Modeling further revealed a conserved, covalent inhibition mechanism involving a P1′ cysteine and offered structural explanations for prime- and non-prime side selectivity, expanding current understanding of legumain’s transpeptidase activity.

Despite these advances, *in silico* screening has limitations. In particular, AlphaFold does not yet account for pH-dependent conformational changes or solvent-mediated interactions, which are likely to influence specificity and catalysis in acidic compartments such as lysosomes or vacuoles. Taken together, this hybrid experimental-computational approach (i) identified sequence features that strongly enhance legumain-mediated peptide cyclization, (ii) developed an optimized cyclization substrate (SFTI_opt_) for hLEG with >60-fold improved activity over the original peptide, (iii) mapped prime- and non-prime side preferences for both human and plant enzymes (Fig. 11), (iv) uncovered a covalent inhibition mechanism via P1′ cysteine, (v) uncovered a *k*_cat_ switch implemented in the substrate, and (vi) demonstrated that computational predictions can guide the design of high-performance substrates and inhibitors across legumain family members and potentially other cysteine proteases.

## EXPERIMENTAL SECTION

### Protein production of human legumain and *Arabidopsis thaliana* legumain isoform β

Human legumain was expressed, purified, and activated as described previously ^19^. Preparation of *A. thaliana* legumain isoform β (AtLEGβ) was carried out as reported earlier ^36^. Briefly, LEXSY P10 cells containing the wild-type human legumain (hLEG), or the V155G-D160Y legumain mutant expression construct or the AtLEGβ expression construct were grown in BHI medium (Jena Bioscience) supplemented with 5 mg/ml porcine Hemin (Jena Bioscience), 50 units/ml penicillin and 50 mg/ml streptomycin (Pen-Strep, Jena Bioscience). Protein expression was carried out in a 500 ml shaking culture (140 rev/min, 26 °C) inoculated 1:10 with transfected strain culture for 2 days. Recombinant protein was removed from the LEXSY supernatant via Ni^2+^ purification using Ni-NTA Superflow resin (Qiagen, Hilden, Germany). Eluates were concentrated using Amicon Ultra centrifugal filter units (10 kDa molecular-weight cutoff, Millipore) and the buffer was exchanged using PD-10 columns (GE Healthcare) to 20 mM Hepes pH 7.0, 50 mM NaCl, 2 mM DTT in case of hLEG or 20 mM Hepes pH 7.2 and 50 mM NaCl in case of AtLEGβ. Activation of human prolegumain was achieved by adjusting the protein concentration to 1 mg/ml and adding 1 M citric acid (pH 3.5) to a final concentration of 100 mM, followed by incubation for > 16 h at 20 °C. Activation of AtLEGβ was carried out under similar conditions, except that 1 M citric acid (pH 4.2) was used and incubation proceeded for 1 h at 30 °C. Activated hLEG was further purified by size-exclusion chromatography on a Superdex S200 10/300 GL column (GE Healthcare) using an ÄKTA FPLC system, yielding protein in 20 mM citric acid pH 4.0, 50 mM NaCl, and 2 mM DTT. Peak fractions were concentrated to ∼10 mg/ml, aliquoted, and stored at -20 °C. Activated AtLEGβ was purified using PD-10 columns (GE Healthcare) to obtain the protein in 20 mM citric acid pH 4.2 and 50 mM NaCl, then concentrated to approximately 1.5 mg/ml.

### Ligation assay using a positional scanning library

To study the influence of specific positions on the ligase activity of hLEG, a positional scan library was used. This library was based on a peptide derived from the sunflower trypsin inhibitor 1 (SFTI-1) as a starting point (G^1^RCTRSIPPICFPN^14^GLA^17^). In this peptide, positions 1, 2, 12, 13, 15, 16, and 17, representing P2’’, P1’’, P3, P2, P1’, P2’, and P3’ respectively, were varied with the 20 proteinogenic amino acids. This resulted in a crude library of 140 peptide variants, synthesized by Biosynth (United Kingdom). Each peptide variant was individually assayed by incubating 60 nM activated hLEG with 500 µM SFTI peptide in cyclization assay buffer (50 mM citric acid, pH 6.0, 100 mM NaCl). Reactions were carried out for 16 h at 37 °C and terminated by addition of 2 % TFA and freezing at −20 °C. Following C18 purification, peptide cyclization was analyzed by mass spectrometry.

For MALDI-ToF mass spectroscopy 20 µL samples were purified and desalted via self-made ZipTips with C18 resin (Empore™ SPE Disks, matrix active group C1, CDS Analytical). Eluates were dried in a speedvac and resuspended in 5 µL of a solution of 25% acetonitrile and 75% TFA (0.1%) utilizing a sonication bath. 5 µL of matrix (α-Cyano-4-hydroxycinnamic acid, Sigma-aldrich) were mixed with the sample and spotted on 2 positions on a steel target (MTP 348 polished steel) using the dried droplet method. Mass spectra were acquired on a Bruker Autoflex Speed MALDI-ToF system operated in positive ionization mode (calibrated mass range 700–3500 m/z, 500 shots per measurement) using FlexControl v3.4 software. Quantification was performed in FlexAnalysis by integrating the peak areas corresponding to the unprocessed precursor, the linearized product, and the cyclic product. The sum of all three peaks represented the total peptide signal at the start of the reaction.

For the positional scanning experiments, peak areas were quantified from single measurements of 500 laser shots. The relative abundance of cyclic product was calculated as the ratio of the cyclic product peak area to the total peptide signal, and values were normalized to those obtained for the peptide containing alanine at the respective variable position. Amino acid substitutions that enhanced cyclization efficiency were identified by plotting normalized cyclic product yields as a heat map. Selected peptides were resynthesized at >95% purity (Thermo Fisher Scientific, Waltham, USA). The variants tested included P2–P13G, P2′–L16K, and the double mutant P13G–L16K. Cyclization of these variants was performed under identical assay conditions, and reaction products were analyzed and quantified as described above. Based on these results, an optimized peptide (SFTI_opt_) was deduced with the consensus sequence G¹–R/K–…–F–G–N¹⁴–G–R/K–X. Experiments that were quantified via mass spectrometry were done with a single replicate (n = 1).

### Kinetic analysis of SFTI cyclisation

Time-course experiments were performed with the optimized substrate SFTI_opt_ to evaluate the reaction rate and stability of hLEG. Predilutions (0.72 µM) of hLEG were incubated for 2 min at 37 °C in cyclisation assay buffer (50 mM citric acid pH 6.0, 100 mM NaCl), or dilution buffer (20 mM citric acid pH 4.0, 100 mM NaCl). This preincubation was immediately followed by incubation of 60 nM of legumain with 500 µM SFTI_opt_ peptide in cyclisation assay buffer (50 mM citric acid pH 6.0, 100 mM NaCl). Samples were taken at specified timepoints (directly after start of the reaction, after 30 sec, and after 1, 2, 5, 10, and 30 min) by stopping the reaction of a 20 µL aliquot with 2% TFA and freezing (-20 °C).

The formation of cyclic product over a broad range of substrate concentrations was monitored to assess binding characteristics of hLEG toward SFTI_opt_. The reaction time was set to 2 min to ensure legumain was still stable in the assay conditions. Serial dilutions of SFTI_opt_ ranging from 1000 µM to 15.5 µM were incubated with 120 nM of activated legumain in cyclisation assay buffer (50 mM citric acid pH 6.0, 100 mM NaCl). After 2 min, the reactions were quenched with 2% TFA and frozen (-20 °C). Subsequently, all samples were desalted via ZipTips and analysed by MALDI-ToF mass spectroscopy as described above.

For kinetic measurements, 10 spectra (500 laser shots each; two sample spots with five measurements per spot, totaling 5000 shots per sample) were acquired and averaged. Enzyme kinetic data for SFTI_opt ligation by hLEG were analyzed in GraphPad Prism (v9.3.1; GraphPad Software, La Jolla, CA) using the built-in Michaelis–Menten and Allosteric sigmoidal models.

The allosteric sigmoidal model was defined by the equation:

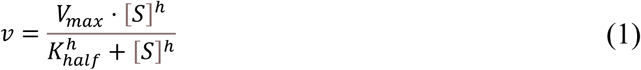

where *v* is the reaction velocity, *V*ₘₐₓ the maximal velocity, [S] the substrate concentration, *h* the Hill slope, and *K*ₕₐₗₓ the substrate concentration at half-maximal velocity. *K*ₕₐₗₓʰ (denoted *K*′) corresponds to the apparent *K_m_*. Standard errors of the curve fits were reported for all fitted parameters.

### Enzyme kinetic assays using fluorescent substrates

K_m_ values for the substrates Z-AAN-AMC (benzyloxycarbonyl-Ala-Ala-Asn-7-amino-4-methylcoumarin) and Z-VAN-AMC (benzyloxycarbonyl-Val-Ala-Asn-7-amino-4-methylcoumarin) toward hLEG were determined in 50 mM citric acid buffer (pH 5.5) containing 100 mM NaCl and 0.05% Tween-20. Reactions were carried out at 37 °C with 2 nM hLEG. Substrate concentrations ranged from 100 µM to 0.787 µM for Z-AAN-AMC and from 75 µM to 0.28 µM for Z-VAN-AMC.

Reactions were initiated by addition of the enzyme, and fluorescence was monitored in a microplate reader (CLARIOstar Plus with Voyager v2405 software, BMG Labtech) at 380 nm excitation and 460 nm emission. All measurements were performed in triplicate (n = 3). Data analysis was performed in GraphPad Prism (v9.3.1; GraphPad Software, La Jolla, CA) using the built-in Michaelis–Menten model for Z-AAN-AMC and the Substrate inhibition model for Z-VAN-AMC. Data points corresponding to 23.73 µM Z-VAN-AMC were excluded from analysis, as a linear fit of the initial slope was not possible. Standard errors of the curve fits are reported for all fitted parameters.

### Peptide competition assays

Inhibition of hLEG and plant legumain (*Arabidopsis thaliana* AtLEGβ) by the peptides GVTN-CR and GVTN-GR was evaluated using a competition assay. Both peptides were synthesized by Thermo Fisher Scientific (Waltham, USA).

Each peptide (2 mM) was preincubated with the fluorogenic legumain substrate Z-AAN-AMC (25 µM for hLEG; 50 µM for AtLEGβ) in activity assay buffer (50 mM citric acid, pH 5.5, 100 mM NaCl, 0.05% Tween-20, 2 mM DTT) for 15 min at 37 °C. Control reactions contained DMSO in place of peptide. Reactions were initiated by adding 2 nM hLEG or 20 nM AtLEGβ. To assess whether inhibition by GVTN-CR was mediated by disulfide bond formation, the assay was repeated using buffer supplemented with 20 mM DTT.

Fluorescence was measured in a microplate reader (CLARIOstar Plus with Voyager v2405 software, BMG Labtech) at 380 nm excitation and 460 nm emission. All measurements were performed in triplicate (*n* = 3). Data were analyzed in GraphPad Prism (v9.3.1; GraphPad Software, La Jolla, CA). Reaction progress curves were fitted, and the slope of each trace was used as a measure of residual legumain activity in the presence of inhibitor peptides.

### AlphaFold Modelling

AlphaFold 3 ^31^ calculations were carried out on two workstation PCs either equipped with Nvidia Geforce RTX 4060 or Nvidia Geforce RTX 4070 Super GPUs. Models of hLEG in complex with peptides derived from the SFTI-based positional scanning assay were prepared. Multiple sequence alignments were enabled, while the use of structural templates was disabled for the peptides. For the *in silico* substrate specificity screening, legumain:peptide structures were calculated using a Python-based script. The first step of the script included the generation of a combinatorial peptide library. In the second step all peptide variants were iteratively applied to structure prediction in complex with the proteases. To save computation time pre-calculated multiple sequence alignments of the protease sequences were used for the AlphaFold 3 calculations. Multiple sequence alignments were in general disabled for the peptides. The use of structural templates was disabled for proteases and for the peptides. For hLEG and *Arabidopsis thaliana* legumain isoform beta (AtLEGβ) the sequences Gly26-Lys289 (uniport accession code Q99538) and Val48-Asn329 (uniport accession code Q39044) were used respectively. The AlphaFold 3 output was analyzed for the ipTM-values, the predicted aligned error (PAE)-values and the predicted Local Distance Difference Test (pLDDT) values of the peptides (average and residue-wise) using another Python-based script. The output was ranked by the ipTM-values and by the average pLDDT values of the peptides. OpenAI GPT-4o and o4-mini high aided with writing of the Python code. Results were visualized using heatmaps colored according to ipTM scores.

## ASSOCIATED CONTENT

### Supporting Information

The following file is available free of charge. Supporting Figures (PDF)

## AUTHOR INFORMATION

### Author Contributions

The manuscript was written through contributions of all authors. All authors have given approval to the final version of the manuscript.

### Notes

The authors declare no competing financial interest.

#### ABBREVIATIONS

AEP: asparaginyl endopeptidase;
SFTI: sunflower trypsin inhibitor;
AtLEGβ: Arabidopsis thaliana legumain β.

## ACKNOWLEDGMENT

The authors wish to thank Martina Wiesbauer and Sabine Ullrich for technical assistance.

