## Supplemental Information for "Positional Scanning and Computational Modeling Reveal Determinants of Legumain Transpeptidase Activity"

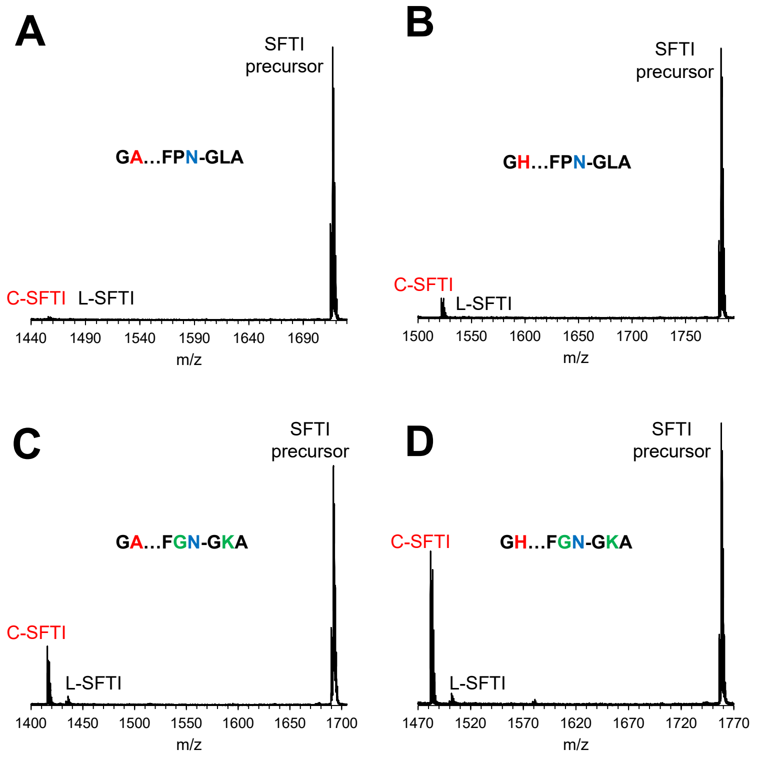


**Supplementary Figure 1. Human legumain prefers positively charged amino acids at position P2"for efficient ligation. (A-D)** MALDI-ToF mass spectra of selected SFTI variants after incubation with human legumain (hLEG): **(A)** $G^{1}A^{2}$…$FPN^{14}GLA$ peptide, **(B)** $G^{1}H^{2}$…$FG^{13}N^{14}GLA$ peptide, **(C)** $G^{1}A^{2}$…$FG^{13}N^{14}GK^{16}A$ peptide, and **(D)** $G^{1}H^{2}$…$FG^{13}N^{14}GK^{16}A$ peptide. The indicated peptides were resynthesized at >95% purity. Peaks corresponding to the unprocessed SFTI precursor, the cyclic SFTI product (C-SFTI), and the linear intermediate lacking residues $G^{15}LA^{17}$ or $G^{15}KA^{17}$ (L-SFTI) are annotated.

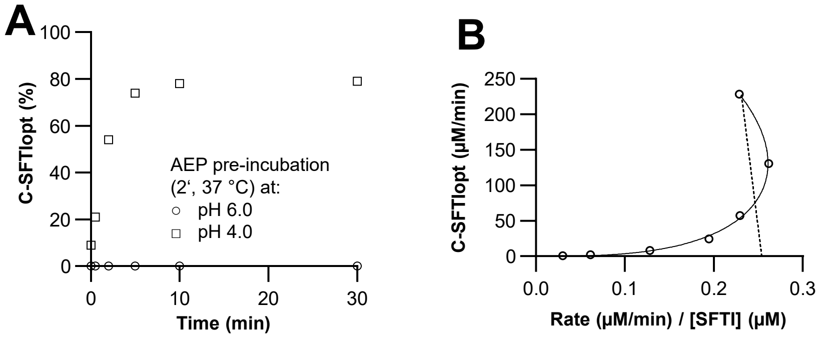


**Supplementary Figure 2. Transpeptidation efficiency of legumain depends on its fold stability and substrate-binding kinetics. (A)** Human legumain was preincubated at pH 4.0 or pH 6.0 for 2 min at 37 °C. Cyclization was subsequently assayed at pH 6.0 with 60 nM enzyme. The relative amount of cyclized $SFTI_{opt}$ (C-$SFTI_{opt}$) was expressed as the percentage of total $SFTI_{opt}$ in the reaction. **(B)** Eadie-Hofstee transformation of human legumain-catalyzed $SFTI_{opt}$ cyclization. Points represent experimental data; the solid line shows the fit to the allosteric sigmoidal model (Eq. 1), and the dashed line shows the fit to the Michaelis-Menten model. The Eadie-Hofstee plot supports that the SFTI-cyclisation reaction does not follow classical Michaelis-Menten kinetics.

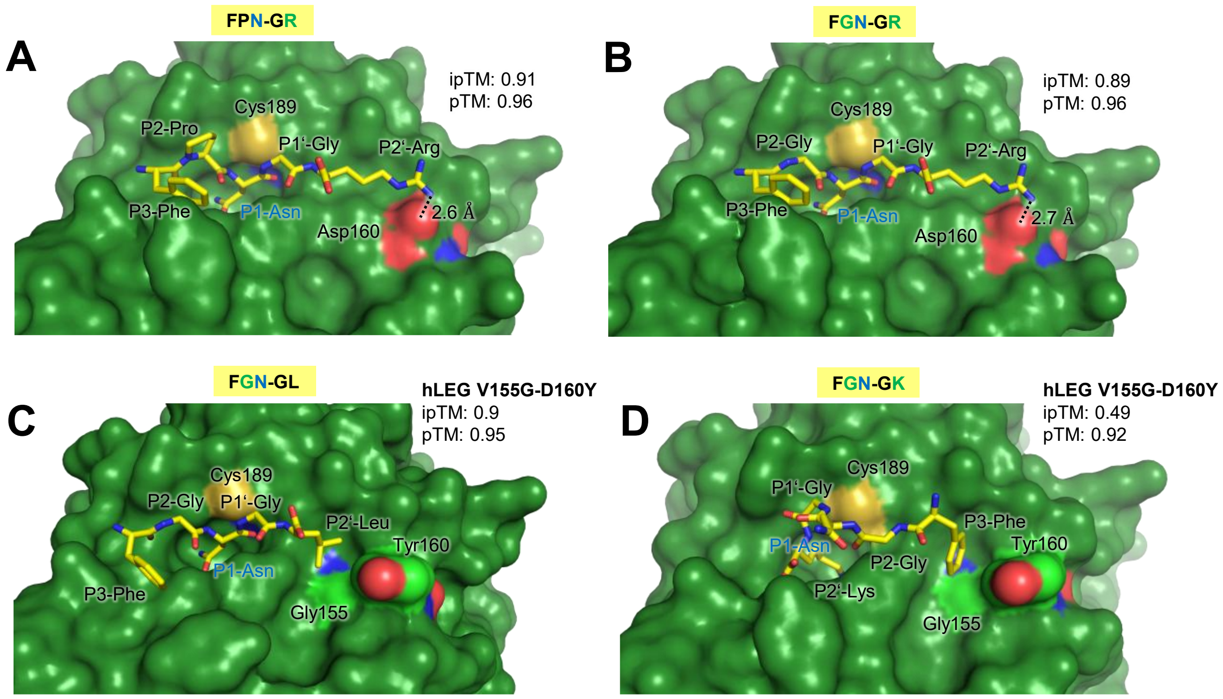


**Supplementary Figure 3. The S2' pocket is a key determinant of substrate specificity. (A,B)** AlphaFold 3 models of human legumain bound to the indicated peptides: **(A)** FPN[14]–GR, **(B)** FGN[14]–GR. **(C, D)** AlphaFold 3 models of human legumain V155G-D160Y bound to the indicated peptides: **(C)** FGN[14]–GL, **(D)** FGN[14]–GK.

| Linear product (normalized, Ala = 1) | | | | | | | |
|---|---|---|---|---|---|---|---|
| | **P1"** | **P2"** | **P3** | **P2** | **P1'** | **P2'** | **P3'** |
| **A** | 1.0 | 1.0 | 1.0 | 1.0 | 1.0 | 1.0 | 1.0 |
| **C** | 0.1 | 0.2 | 0.3 | 1.8 | 7.6 | 0.9 | 5.7 |
| **D** | 0.5 | 0.9 | 0.1 | 0.2 | 0.2 | 0.6 | 1.5 |
| **E** | 0.8 | 0.1 | 0.1 | 1.3 | 0.5 | 0.4 | 0.8 |
| **F** | 0.4 | 0.3 | 0.3 | 0.1 | 0.6 | 0.1 | 0.0 |
| **G** | 0.4 | 0.7 | 0.1 | 0.2 | 5.6 | 0.4 | 1.8 |
| **H** | 0.5 | 0.6 | 0.1 | 0.2 | 5.9 | 1.7 | 2.2 |
| **I** | 0.8 | 0.4 | 0.1 | 0.1 | 0.3 | 0.7 | 0.3 |
| **K** | 0.7 | 1.4 | 0.1 | 0.2 | 3.0 | 2.9 | 3.2 |
| **L** | 0.5 | 0.3 | 0.2 | 0.1 | 0.4 | 0.4 | 2.7 |
| **M** | 0.3 | 0.4 | 0.3 | 0.2 | 0.5 | 0.3 | 0.3 |
| **N** | 0.2 | 0.7 | 0.3 | 0.2 | 0.5 | 0.3 | 0.7 |
| **P** | 0.3 | 0.2 | 0.1 | 0.3 | 0.9 | 0.3 | 0.2 |
| **Q** | 0.0 | 0.1 | 0.1 | 0.6 | 0.7 | 0.6 | 0.4 |
| **R** | 0.9 | 0.7 | 0.9 | 1.4 | 1.7 | 2.3 | 0.9 |
| **S** | 0.2 | 0.1 | 0.7 | 1.0 | 17.0 | 0.8 | 1.1 |
| **T** | 0.2 | 0.1 | 0.2 | 0.2 | 0.7 | 1.3 | 2.3 |
| **V** | 0.8 | 0.1 | 0.2 | 0.7 | 1.3 | 0.2 | 0.3 |
| **W** | 0.0 | 0.1 | 0.1 | 0.2 | | 0.0 | 0.3 |
| **Y** | 0.4 | 0.2 | 0.1 | 0.1 | 0.6 | 0.4 | 0.7 |

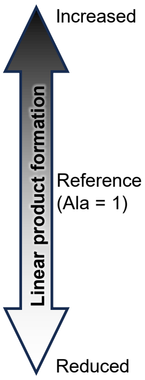


**Supplementary Figure 4. Legumain hydrolase activity analyzed based on the amount of linear product formed.** Results of the positional peptide scanning assay using a crude peptide library of 140 variants are summarized in a heatmap. The relative amount of linear SFTI product (L-SFTI) formed was normalized to the peptide containing alanine at the respective position.

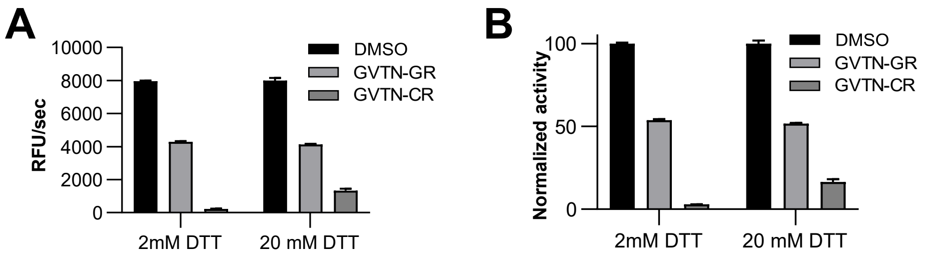


**Supplementary Figure 5. P1′-Cys interacts with human legumain through disulfide bond formation. (A)** Competition assays of human legumain with the peptides GVTN-CR and GVTN-GR. Each peptide (2 mM) was preincubated with the fluorogenic substrate Z-AAN-AMC in assay buffer (pH 5.5) containing either 2 mM or 20 mM DTT. Reactions were initiated by the addition of 2 nM human legumain, and fluorescence was monitored over 20 min at 37 °C. **(B)** Same experiment as in **(A)**, with data normalized to the respective DMSO control reactions.

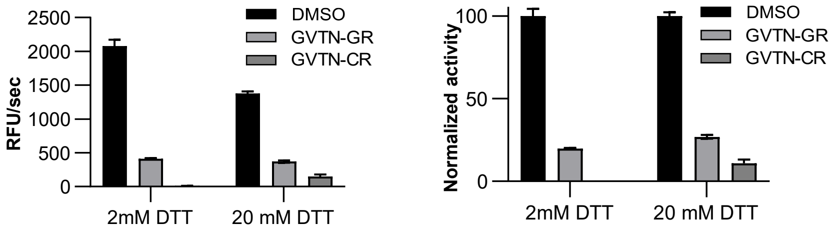


**Supplementary Figure 6. P1′-Cys interacts with *Arabidopsis thaliana* legumain β (AtLEGβ) through disulfide bond formation.** **(A)** Competition assays of AtLEGβ with the peptides GVTN-CR and GVTN-GR. Each peptide (2 mM) was preincubated with the fluorogenic substrate Z-AAN-AMC in assay buffer (pH 5.5) containing either 2 mM or 20 mM DTT. Reactions were initiated by the addition of 20 nM AtLEGβ, and fluorescence was monitored over 20 min at 37 °C. (**B**) Same experiment as in **(A)**, with data normalized to the respective DMSO control reactions.

**A**

**AtLEGβ + AXXN-CL**
(colored according to ipTM score)

| P2 \ P3 | A | C | D | E | F | G | H | I | K | L | M | N | P | Q | R | S | T | V | W | Y |
|---|---|---|---|---|---|---|---|---|---|---|---|---|---|---|---|---|---|---|---|---|
| A | 0.94 | 0.91 | 0.93 | 0.94 | 0.92 | 0.93 | 0.94 | 0.94 | 0.93 | 0.93 | 0.94 | 0.94 | 0.94 | 0.94 | 0.93 | 0.93 | 0.95 | 0.95 | 0.94 | 0.93 |
| C | 0.94 | 0.91 | 0.93 | 0.94 | 0.93 | 0.93 | 0.94 | 0.94 | 0.93 | 0.93 | 0.94 | 0.94 | 0.94 | 0.94 | 0.93 | 0.94 | 0.95 | 0.94 | 0.93 | 0.92 |
| D | 0.94 | 0.87 | 0.93 | 0.94 | 0.92 | 0.93 | 0.93 | 0.94 | 0.93 | 0.94 | 0.93 | 0.94 | 0.93 | 0.94 | 0.94 | 0.93 | 0.95 | 0.95 | 0.94 | 0.93 |
| E | 0.94 | 0.90 | 0.92 | 0.94 | 0.92 | 0.93 | 0.93 | 0.95 | 0.92 | 0.93 | 0.92 | 0.93 | 0.93 | 0.94 | 0.93 | 0.93 | 0.95 | 0.95 | 0.93 | 0.93 |
| F | 0.94 | 0.82 | 0.93 | 0.93 | 0.92 | 0.92 | 0.92 | 0.93 | 0.91 | 0.92 | 0.91 | 0.93 | 0.92 | 0.93 | 0.93 | 0.92 | 0.94 | 0.94 | 0.93 | 0.92 |
| G | 0.92 | 0.87 | 0.93 | 0.94 | 0.93 | 0.90 | 0.94 | 0.94 | 0.92 | 0.92 | 0.93 | 0.93 | 0.93 | 0.93 | 0.93 | 0.92 | 0.94 | 0.95 | 0.93 | 0.93 |
| H | 0.94 | 0.88 | 0.93 | 0.94 | 0.92 | 0.93 | 0.94 | 0.94 | 0.93 | 0.93 | 0.92 | 0.94 | 0.93 | 0.93 | 0.93 | 0.93 | 0.94 | 0.94 | 0.93 | 0.93 |
| I | 0.94 | 0.82 | 0.94 | 0.94 | 0.93 | 0.94 | 0.92 | 0.94 | 0.92 | 0.92 | 0.93 | 0.93 | 0.92 | 0.94 | 0.93 | 0.93 | 0.95 | 0.95 | 0.93 | 0.92 |
| K | 0.94 | 0.91 | 0.94 | 0.94 | 0.92 | 0.94 | 0.94 | 0.94 | 0.93 | 0.93 | 0.93 | 0.94 | 0.94 | 0.93 | 0.94 | 0.93 | 0.95 | 0.95 | 0.94 | 0.93 |
| L | 0.94 | 0.87 | 0.93 | 0.93 | 0.91 | 0.94 | 0.92 | 0.93 | 0.92 | 0.92 | 0.93 | 0.94 | 0.92 | 0.94 | 0.93 | 0.93 | 0.94 | 0.95 | 0.93 | 0.92 |
| M | 0.94 | 0.91 | 0.93 | 0.94 | 0.93 | 0.94 | 0.94 | 0.94 | 0.93 | 0.93 | 0.93 | 0.94 | 0.94 | 0.94 | 0.94 | 0.93 | 0.95 | 0.95 | 0.93 | 0.92 |
| N | 0.94 | 0.91 | 0.93 | 0.94 | 0.92 | 0.93 | 0.94 | 0.94 | 0.93 | 0.93 | 0.94 | 0.94 | 0.93 | 0.94 | 0.94 | 0.93 | 0.95 | 0.95 | 0.93 | 0.93 |
| P | 0.94 | 0.90 | 0.94 | 0.94 | 0.93 | 0.89 | 0.94 | 0.94 | 0.93 | 0.93 | 0.94 | 0.93 | 0.94 | 0.94 | 0.93 | 0.93 | 0.95 | 0.95 | 0.94 | 0.94 |
| Q | 0.94 | 0.90 | 0.93 | 0.94 | 0.92 | 0.93 | 0.93 | 0.94 | 0.93 | 0.93 | 0.93 | 0.94 | 0.94 | 0.93 | 0.93 | 0.93 | 0.95 | 0.95 | 0.94 | 0.93 |
| R | 0.94 | 0.92 | 0.93 | 0.94 | 0.92 | 0.94 | 0.94 | 0.94 | 0.93 | 0.93 | 0.93 | 0.94 | 0.94 | 0.94 | 0.94 | 0.93 | 0.95 | 0.95 | 0.93 | 0.92 |
| S | 0.93 | 0.86 | 0.93 | 0.94 | 0.92 | 0.93 | 0.94 | 0.95 | 0.93 | 0.93 | 0.94 | 0.93 | 0.94 | 0.93 | 0.93 | 0.93 | 0.95 | 0.95 | 0.94 | 0.92 |
| T | 0.94 | 0.90 | 0.94 | 0.94 | 0.92 | 0.93 | 0.94 | 0.95 | 0.94 | 0.93 | 0.94 | 0.94 | 0.94 | 0.94 | 0.94 | 0.94 | 0.95 | 0.95 | 0.93 | 0.92 |
| V | 0.94 | 0.88 | 0.94 | 0.94 | 0.92 | 0.94 | 0.92 | 0.95 | 0.93 | 0.93 | 0.93 | 0.94 | 0.93 | 0.94 | 0.93 | 0.94 | 0.95 | 0.95 | 0.93 | 0.92 |
| W | 0.93 | 0.82 | 0.93 | 0.93 | 0.92 | 0.91 | 0.93 | 0.93 | 0.92 | 0.92 | 0.92 | 0.93 | 0.92 | 0.93 | 0.93 | 0.92 | 0.94 | 0.94 | 0.93 | 0.93 |
| Y | 0.94 | 0.86 | 0.92 | 0.93 | 0.92 | 0.93 | 0.93 | 0.93 | 0.93 | 0.92 | 0.92 | 0.93 | 0.92 | 0.93 | 0.93 | 0.92 | 0.93 | 0.94 | 0.92 | 0.92 |

High ipTM
ipTM (AtLEGβ + peptide)
Low ipTM

**B**

**AtLEGβ + XXAN-CL**
(colored according to ipTM score)

| P3 \ P4 | A | C | D | E | F | G | H | I | K | L | M | N | P | Q | R | S | T | V | W | Y |
|---|---|---|---|---|---|---|---|---|---|---|---|---|---|---|---|---|---|---|---|---|
| A | 0.94 | 0.90 | 0.93 | 0.93 | 0.94 | 0.94 | 0.95 | 0.95 | 0.94 | 0.95 | 0.94 | 0.94 | 0.94 | 0.94 | 0.94 | 0.93 | 0.94 | 0.94 | 0.94 | 0.94 |
| C | 0.91 | 0.92 | 0.91 | 0.91 | 0.94 | 0.89 | 0.92 | 0.92 | 0.90 | 0.93 | 0.90 | 0.88 | 0.89 | 0.91 | 0.87 | 0.89 | 0.91 | 0.93 | 0.92 | 0.93 |
| D | 0.93 | 0.92 | 0.92 | 0.92 | 0.95 | 0.94 | 0.94 | 0.94 | 0.93 | 0.95 | 0.93 | 0.93 | 0.93 | 0.93 | 0.93 | 0.92 | 0.94 | 0.94 | 0.93 | 0.95 |
| E | 0.94 | 0.92 | 0.92 | 0.94 | 0.94 | 0.94 | 0.95 | 0.95 | 0.95 | 0.95 | 0.94 | 0.94 | 0.94 | 0.93 | 0.94 | 0.94 | 0.95 | 0.95 | 0.93 | 0.95 |
| F | 0.92 | 0.91 | 0.93 | 0.92 | 0.93 | 0.93 | 0.92 | 0.93 | 0.92 | 0.94 | 0.93 | 0.93 | 0.92 | 0.93 | 0.93 | 0.93 | 0.93 | 0.93 | 0.93 | 0.93 |
| G | 0.93 | 0.88 | 0.93 | 0.93 | 0.94 | 0.94 | 0.94 | 0.93 | 0.92 | 0.93 | 0.93 | 0.93 | 0.94 | 0.93 | 0.92 | 0.93 | 0.94 | 0.93 | 0.94 | 0.94 |
| H | 0.94 | 0.92 | 0.94 | 0.93 | 0.94 | 0.94 | 0.94 | 0.94 | 0.93 | 0.94 | 0.94 | 0.94 | 0.94 | 0.93 | 0.93 | 0.93 | 0.94 | 0.94 | 0.94 | 0.94 |
| I | 0.94 | 0.91 | 0.94 | 0.94 | 0.95 | 0.94 | 0.96 | 0.95 | 0.95 | 0.95 | 0.95 | 0.93 | 0.94 | 0.93 | 0.94 | 0.94 | 0.95 | 0.95 | 0.95 | 0.94 |
| K | 0.93 | 0.89 | 0.93 | 0.93 | 0.93 | 0.93 | 0.94 | 0.94 | 0.94 | 0.94 | 0.93 | 0.92 | 0.93 | 0.93 | 0.93 | 0.92 | 0.93 | 0.94 | 0.93 | 0.94 |
| L | 0.93 | 0.89 | 0.93 | 0.93 | 0.94 | 0.93 | 0.93 | 0.94 | 0.92 | 0.94 | 0.93 | 0.92 | 0.93 | 0.92 | 0.92 | 0.93 | 0.93 | 0.93 | 0.93 | 0.93 |
| M | 0.94 | 0.90 | 0.93 | 0.92 | 0.95 | 0.94 | 0.94 | 0.94 | 0.94 | 0.95 | 0.94 | 0.92 | 0.94 | 0.92 | 0.94 | 0.93 | 0.93 | 0.94 | 0.94 | 0.94 |
| N | 0.94 | 0.90 | 0.93 | 0.93 | 0.95 | 0.94 | 0.93 | 0.94 | 0.93 | 0.93 | 0.94 | 0.93 | 0.94 | 0.93 | 0.94 | 0.94 | 0.94 | 0.94 | 0.94 | 0.94 |
| P | 0.94 | 0.90 | 0.93 | 0.92 | 0.93 | 0.93 | 0.94 | 0.95 | 0.91 | 0.94 | 0.93 | 0.93 | 0.95 | 0.93 | 0.93 | 0.92 | 0.94 | 0.95 | 0.94 | 0.94 |
| Q | 0.94 | 0.92 | 0.92 | 0.93 | 0.94 | 0.94 | 0.95 | 0.95 | 0.94 | 0.95 | 0.94 | 0.93 | 0.93 | 0.93 | 0.94 | 0.93 | 0.94 | 0.95 | 0.93 | 0.95 |
| R | 0.93 | 0.90 | 0.93 | 0.93 | 0.94 | 0.93 | 0.93 | 0.94 | 0.92 | 0.94 | 0.93 | 0.93 | 0.93 | 0.93 | 0.93 | 0.93 | 0.93 | 0.94 | 0.93 | 0.94 |
| S | 0.93 | 0.91 | 0.93 | 0.93 | 0.94 | 0.94 | 0.94 | 0.94 | 0.93 | 0.94 | 0.93 | 0.93 | 0.93 | 0.93 | 0.93 | 0.93 | 0.94 | 0.94 | 0.93 | 0.94 |
| T | 0.95 | 0.92 | 0.93 | 0.95 | 0.96 | 0.95 | 0.96 | 0.95 | 0.95 | 0.96 | 0.95 | 0.95 | 0.95 | 0.92 | 0.96 | 0.94 | 0.95 | 0.95 | 0.96 | 0.96 |
| V | 0.95 | 0.92 | 0.94 | 0.94 | 0.95 | 0.95 | 0.96 | 0.95 | 0.95 | 0.95 | 0.95 | 0.95 | 0.95 | 0.94 | 0.95 | 0.94 | 0.95 | 0.95 | 0.95 | 0.95 |
| W | 0.94 | 0.92 | 0.94 | 0.93 | 0.96 | 0.94 | 0.93 | 0.94 | 0.93 | 0.95 | 0.95 | 0.94 | 0.94 | 0.93 | 0.94 | 0.94 | 0.94 | 0.94 | 0.94 | 0.95 |
| Y | 0.93 | 0.91 | 0.93 | 0.93 | 0.94 | 0.94 | 0.93 | 0.93 | 0.93 | 0.94 | 0.94 | 0.94 | 0.93 | 0.93 | 0.94 | 0.93 | 0.94 | 0.93 | 0.94 | 0.94 |

High ipTM
ipTM (AtLEGβ + peptide)
Low ipTM

**Supplementary Figure 6. Substrate specificity is relatively broad on the non-prime side of *Arabidopsis thaliana* legumain β (AtLEGβ).** Results of the AlphaFold 3-based *in silico* substrate specificity screening using the sequence of AtLEGβ and the XXXN-CL peptide are shown. To enable two-dimensional visualization, either the P2 position was fixed as alanine **(A)**

or the P4 position was fixed as alanine **(B)**. Heatmaps are colored according to the ipTM score of the respective legumain-substrate complexes.
